# A chromosome-scale and haplotype-resolved genome assembly of tetraploid blackberry (*Rubus* L. subgenus *Rubus* Watson)

**DOI:** 10.1101/2023.04.24.538158

**Authors:** Dev Paudel, Ze Peng, Saroj Parajuli, S. Brooks Parrish, Zhanao Deng

## Abstract

**Background:** Blackberries (*Rubus* subgenus *Rubus*) are a major berry crop consumed globally for being a rich source of anthocyanins and antioxidants, and their unique flavor. However, breeding for fruit improvement in blackberry has been significantly hindered by the scarcity of genomic resources and the genetic complexity of traits. The blackberry genome has been particularly challenging to assemble, largely due to its polyploid nature.

**Findings:** We present the first chromosome-scale and haplotype-phased genome assembly for the cultivated primocane-fruiting, thornless tetraploid blackberry selection BL1 (*Rubus* L. subgenus *Rubus* Watson). The tetraploid genome assembly was generated using the Oxford Nanopore Technology (ONT) and Hi-C scaffolding, comprising 919 Mb placed on 27 pseudochromosomes with an N50 of 35.73 Mb. The assembly covers >92% of the genome length and contains over 98% of complete BUSCOs. Repetitive sequences constitute 57% of the assembly, with the long terminal repeats (LTR) being the most abundant class. A total of 87,968 protein-coding genes were predicted, of which, 82% were functionally annotated. Gene expression analyses identified candidate genes and transcription factors related to thornlessness in blackberries, including MYB16, lysine histidine transporters, PDR1, Caffeoyl-CoA, glycosylphosphatidylinositol-anchored lipid protein transfer 1, DRN, beta-ketoacyl reductase, and homocysteine S-methyltransferase 3.

**Conclusions:** The utility of this genome has been demonstrated in this study by identifying candidate genes related to thornlessness in blackberry. Resequencing of tetraploid blackberry cultivars/selections with different horticultural characteristics revealed genes that could impact fruiting habit and disease resistance/susceptibility. This tetraploid reference genome will serve as a valuable resource to accelerate genetic analysis and breeding of this important berry crop, enabling the development of improved varieties with enhanced traits.

## 1 Introduction

Blackberries (*Rubus* spp.), belonging to the genus *Rubus* subgenus *Rubus* (formerly subgenus *Eubatus*) within the Rosaceae family (Clark et al., 2007), are characterized by their dark purple to deep black color, compound structure, and a combination of juicy, tart, and sweet flavors. They are an exceptional source of anthocyanins, antioxidants, and dietary fibers, offering significant health benefits to consumers. Over the past two decades, a sharp increase in consumer demand has led to substantial expansion of the market for fresh and processed blackberries in the United States (U.S.) and other countries worldwide (Foster et al., 2019). As the fourth most economically important berry crop in the U.S., the country produced 16,850 metric tons (MT) of processed and 1,360 MT of fresh blackberries in 2017 (Morgan, 2022). In 2021, the U.S. imported 122,873 MT of fresh blackberries and 16,738 MT of frozen blackberries, valued at $519 million and $43 million, respectively (USDA, 2021). The global production of blackberries is estimated to be over 900,000 MT, making it a substantial contributor to the international berry industry. The ongoing development and introduction of new, improved cultivars has been instrumental in addressing consumer demands and increasing blackberry production across the globe.

Blackberries possess a number of interesting biological characteristics that distinguish them from other plants, including the biennial growth habit, thorny canes, compound fruit structure, wide adaptation, and high hybridization potential with other species in the *Rubus* genus, which make blackberry species very interesting for studying the biology of these characteristics (Carter et al., 2019; Gustafsson 1942). Blackberry species exhibit varying ploidy levels and chromosome numbers, ranging from diploids (2*n* = 2*x* = 14) to duodecaploid (2*n* = 12*x* = 84) (Thompson, 1997), which allows for significant genetic diversity within these species.

A primary goal of blackberry breeding is to develop high-yielding, thornless cultivars that produce berries with superior qualities and sweet flavors (Du and Qian, 2010). Shoots or canes of blackberries are covered from base to tip with thorny protrusions. Thorns on blackberry leaves and shoots cause damage to berries, increase disease incidences, and reduce marketable yield. Thorns also make it challenging to harvest berries, prune plants, and present hazards to farm workers. Thorns that contaminate machine-harvested fruit can extremely difficult to remove and thorns that end up in the product are unappealing and dangerous (Strik and Buller, 2002). Thornlessness is essential for new blackberry cultivars to be widely adopted and grown by growers. Consequently, understanding the genetic control of blackberry thorns has been an important area of study in blackberry genetics.

A major breakthrough in blackberry breeding in recent years is the development and release of primocane-fruiting cultivars. While the majority of blackberries bear fruit on second-year canes, known as floricanes, primocane-fruiting blackberries can produce fruit on first-year canes. This trait offers remarkable advantages, including the potential for an extended harvest season, increased yield, and the ability to grow in regions with shorter growing seasons or colder climates (Clark et al., 2007). Consequently, this character has the potential to revolutionize blackberry cultivation, significantly benefiting the global berry industry and consumers. This trait was first discovered in the 1940s in wild blackberry (Clark et al., 2007). Decades of breeding and selection efforts led to the release of the first commercial primocane-fruiting blackberry cultivar, ‘Prime-Ark® 45’, in 2009 (Clark and Perkins-Veazie, 2011). Recent research has started to shed light on the genetic control of this trait (Lopez-Medina et al., 2000). However, a more comprehensive understanding of the genetic control and expression of this game-changing trait is still needed. The development of molecular markers or genomic selection tools would greatly assist in incorporating this trait into a wider range of cultivars, further expanding the benefits of primocane-fruiting blackberries.

In commercial cultivation, blackberries are susceptible to several diseases that require regular chemical sprays to maintain healthy plants and crops (Jennings et al., 1991; Hall, 2011; Marin et al., 2022, 2023). Overreliance on chemical treatments not only increases production costs but also lead to negative environmental consequences, making the development of disease-resistant cultivars a high priority. Selection for disease resistance is still in its infancy in blackberries, and research into disease resistance genes (*R* genes) promise to significantly contribute to enhancing blackberry resistance to diseases. By identifying and selecting *R* genes, breeders can develop cultivars with inherent resistance to pathogens, reducing the need for pesticides and improving the sustainability of blackberry cultivation. This development is particularly important for the expansion of blackberry cultivation into subtropical regions, which presents new changes due to increased disease pressures in these climates (Foster et al., 2019).

Despite their economic importance and health benefits, genomic resources for blackberries remain very limited, impeding the utilization of molecular markers and genomic selection in breeding efforts. Within the *Rubus* genus, the genome sequences for the subgenus *Ideaobatus* include diploid black raspberry (*R. occidentalis*) ORUS 4114-3 (VanBuren et al., 2016; VanBuren et al., 2018), red raspberry (*R. idaeus*) cultivar ‘Anitra’ (Davik et al., 2022), and *R. chingii* (Wang et al., 2021). For the subgenus *Rubus*, the genome of diploid *R. argutus* ‘Hillquist’ (Brůna et al., 2023) was recently released. However, publicly available genome sequence data for tetraploid blackberries of subgenus *Rubus* are still lacking. Polyploid genomes, which have multiple copies of each homologous chromosomes, are more complex and challenging to assemble and decipher compared to their diploid counterparts. Therefore, it is crucial to develop high-quality genome assembly for tetraploid blackberries to facilitate research and breeding advancements.

In this study, we present the first chromosome-length, hyplotype-resolved genome assembly and annotation of tetraploid blackberry, providing a valuable resource for facilitating the genetic improvement of this crop through traditional or precision breeding technologies. Using this genome, we explored global haplotype differences among representative blackberry cultivars/selections with different horticultural characteristics and identified candidate genes associated with disease resistance/ susceptibility, thornlessness, and primocane fruiting in blackberry. These findings, along with the reference genome, will serve as a critical resource for discovery and analysis of genetics behind important traits in blackberries, ultimately enabling and streamlining molecular breeding efforts in this important berry crop.

## 2 Results and discussion

### 2.1 Genome sequencing, assembly, and annotation

A phased, tetraploid reference genome for *Rubus* subgenus *Rubus* Watson BL1 selection (BL1) was de novo assembled using a combination of Oxford Nanopore sequencing (811× coverage of the monoploid genome, 160× coverage of the tetraploid genome) and Illumina sequencing (118× coverage of the monoploid genome) to produce highly contiguous pseudochromosomes (Figure 1, Table 1). Utilizing the Hi-C chromosome conformation capture, 95% of contig sequences were organized onto the 27 chromosomes for the four haplotypes. The total length of the final assembly was 919,165,722 bases distributed across 27 chromosome-level pseudomolecules (291,667,923; 262,227,137; 196,366,124; and 125,536,233 for haplotypes A, B, C, and D, respectively). The chromosomes were numbered sequentially based on their synteny to the diploid blackberry Hillquist genome; and the subgenomes were labelled A-D based on their length (Supplementary Figure S1, Supplementary Figure S2). High collinearity was observed among the homoeologous chromosomes (Supplementary Figure S3). The largest scaffold was 55.81 Mb (Chr A6), and the overall N50 was 35.73 Mb, corresponding to the median chromosome length (Table 1). The estimated GC content was 36.97%. Evaluation of the genome’s completeness using 2,326 eukaryotic genes in the BUSCO OrthoDB10 eudicot dataset showed that the BL1 genome was 98.1% complete in gene space. Telomeric repeats were identified towards one of the distal ends of 17 chromosomes, while telomeric repeats were identified at both ends of two chromosomes (A4 and B7) (Supplementary Figure S4). The high completeness, large N50 length, and identification of telomeres indicate high quality of this genome assembly.

**Figure 1.**
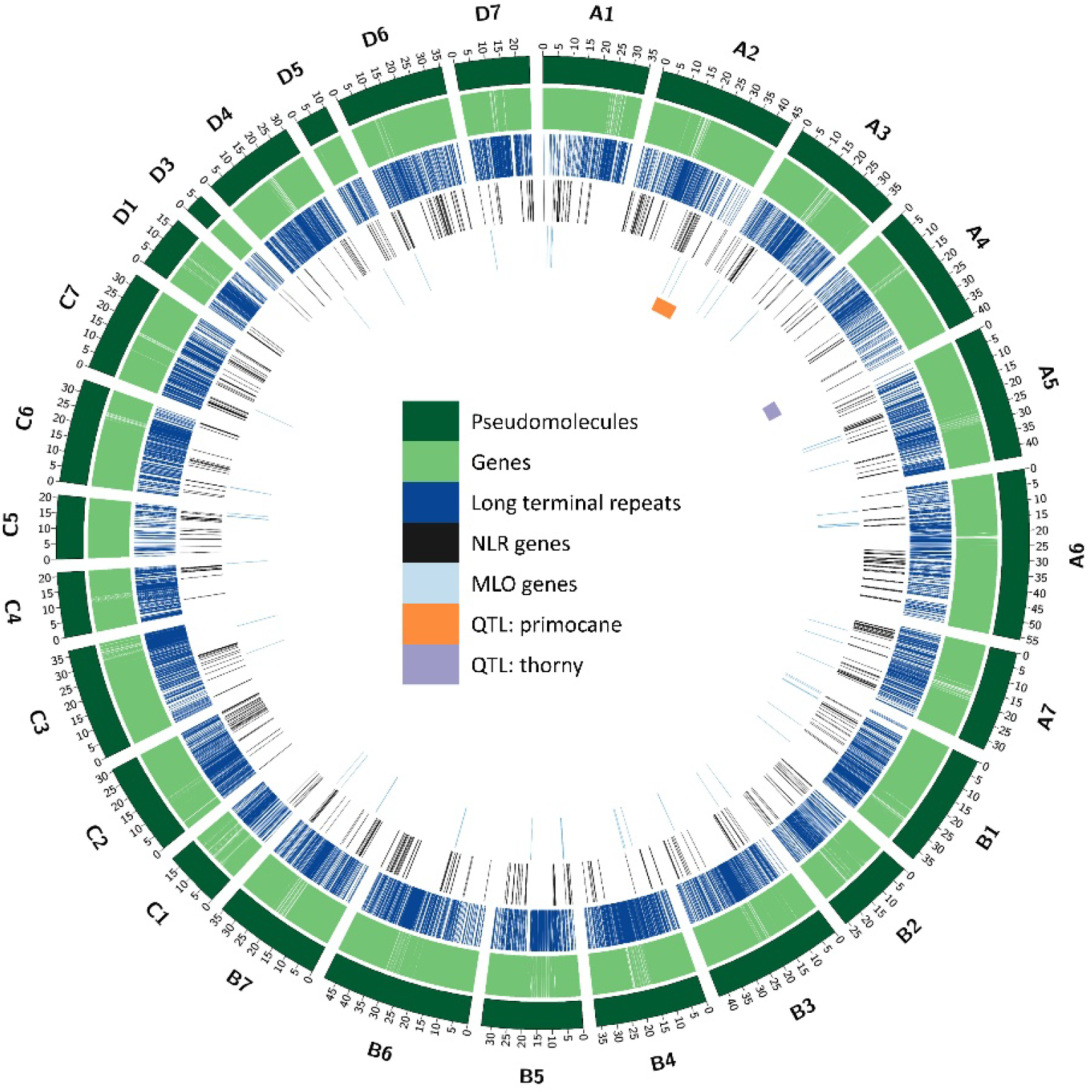
Circos plot showing the characteristics of the *Rubus* L. subgenus *Rubus* Watson BL1 genome. Concentric circles from outside to inside show the following: 1) 27 assembled pseudomolecules (in Mb) (dark green); 2) Locations of predicted gene models (light green); 3) Locations of predicted long terminal repeat (LTR) transposable elements (TEs) (dark blue); 4) Locations of predicted nucleotide-binding site leucine-rich repeat class of disease resistance genes (NLR) (black); 5) Locations of predicted disease susceptibility *Mildew Locus O* (*MLO*) genes (light blue); and 6) Locations of the loci genetically mapped for primocane fruiting (orange) and thornlessness (purple).

**Table 1.** Summary statistics of the BL1 genome assembly and annotation.

|  |  |
| --- | --- |
| Total size of genome assembly | 919.16 Mb |
| Chromosome number ($2n$) | 28 |
| Number of chromosomal pseudomolecules | 27 |
| Longest scaffold | 55.81 Mb |
| Scaffold N50 length | 35.73 Mb |
| Number of unscaffolded contigs | 328 |
| Size of unscaffolded contigs | 42.36 Mb |
| Contig N50 length | 241 Kb |
| GC content | 36.97% |
| Number of gene models | 87,968 |
| Number of genes with isoforms | 6,314 |
| Mean transcript length | 2,179 bp |
| Mean coding sequence length | 1,084 bp |
| Mean exons per transcript | 4.2 |
| Number of transcripts | 94,936 |
| Repetitive elements | 57.44% |

Chromosome staining confirmed 28 chromosomes in the BL1 root tip somatic cells (Supplementary Figure S5). Flow cytometry analysis of BL1 leaf cells revealed that its holoploid nuclear DNA content (2C) was 1.46 ± 0.05 pg, which corresponds to the monoploid 1C_x_ genome size of 0.365 pg (356.97Mb) (Supplementary Figure S6). This estimate is similar to the genome size of diploid *R. argutus* ‘Hillquist’ (337.4 Mb, 1C = 0.345 pg) (Brůna et al., 2023). The 2C nuclear DNA content of diploid *Rubus* species from the *Rubus* subgenus ranges from 0.59 pg (*R. hispidus* and *R. canadensis*) to 0.75 pg (*R. sanctus*) (Meng and Finn, 2002).

The *k*-mer analysis with GenomeScope 2.0 estimated the monoploid genome size of BL1 at 247 Mb, based on the tetraploid model. The calculated *aaab* value was greater than the *aabb* value, suggesting an autopolyploid nature for BL1 (Supplementary Figure S7). This nature was corroborated by the smudgeplot with a bright smudge at AAAB (0.52) (Supplementary Figure S8).

In blackberries, the first genetic linkage map was constructed using 119 simple sequence repeat (SSR) markers (Castro et al., 2013). This number of markers was sufficient for constructing a framework of the blackberry genome. Recently, a maternal haplotype map consisting of 30 linkage groups was developed for blackberry using 2,935 single nucleotide polymorphism (SNP) markers (Brůna et al., 2023). We mapped these SNP markers to the BL1 genome assembly. Most of the linkage groups displayed a one-to-one relationship with the pseudo chromosomes in the BL1 genome (Supplementary Figure S9). However, several pseudochromosomes had fewer numbers of SNPs that were genetically mapped, which might be due to differences in the genome architecture or low numbers of SNPs in the linkage groups.

Currently, only the genome of one diploid blackberry ‘Hillquist’ has been published (Brůna et al., 2023). Compared to the Hillquist assembly (N50 = 38.6 Mb, and the maximum scaffold length = 45.5 Mb), this tetraploid genome assembly has a similar N50 of 35.75 Mb and a larger maximum length of 55.81 Mb (Chromosome A6) (Supplementary Table S1). We were able to incorporate 95% of the contig sequences into 27 chromosomes for the four haplotypes of the tetraploid blackberry, compared to seven collapsed chromosomes in the Hillquist genome. Based on the estimated genome size through *k*-mer analysis, the BL1 genome covers more than 92% of the genome length and contains over 98% of complete BUSCOs (Supplementary Table S2), which indicates a high level of completeness. In the BL1 genome assembly, repetitive elements account for 57.44% of the genome, slightly higher than the repeat content in the Hillquist genome (52.8%). Several chromosomal rearrangements are visible in the BL1 genome when compared to the Hillquist genome. However, further investigation is necessary to confirm these rearrangements.

Inherent polyploidy in tetraploid blackberries makes it challenging to assemble and accurately phase the genomes. While using Nanopore with Hi-C has enabled us to phase most of the genome (>90%), we were still unable to assemble one chromosome (Chr D2), which most likely collapsed during the assembly phases. This collapse is likely due to the chromosome being highly similar to the other homologous chromosomes, causing the assembler to treat the chromosome as if it were one of the others. This chromosome appears to be significant from the perspective of disease resistance, as it contains the highest number of nucleotide-binding site and leucine-rich repeat containing (NLR) genes (see below). Consequently, it is likely that this chromosome was highly conserved, leading to its collapse in the assembly.

### 2.2 Genome annotation

The BL1 genome contains 57.44% repetitive elements (Supplementary Table S3). Long terminal repeat (LTR) transposable elements are the most predominant class (30.07%) while other repetitive elements are present in much smaller proportions, including DNA elements (1.88%), LINEs (0.96%), and SINEs (0.01%). An integrated strategy combining *ab initio* and the homology-based methods was employed to predict gene models in the genome. We leveraged RNA-Seq data from public and private databases and proteins from the Hillquist genome to make accurate predictions. In total, we identified 87,968 protein-coding genes with 94,936 transcripts (Table 1). The mean gene length is 2,179 bp, and the mean coding sequence length is 1,515 bp, with an average of 4.2 exons per transcript. The total gene length (206,900,785 bp) accounts for 22.5% of the BL1 genome. A total of 6,314 genes have spliced isoforms, with an average of 2.1 isoforms per gene. The total gene number in BL1 is 87,968, compared to 38,503 protein-coding genes in the Hillquist genome assembly. Functional annotation was assigned to 82% of the genes identified in the genome.

### 2.3 Comparative genomics within the Rosaceae

The gene family assignment using existing annotations for the 15 published Rosaceae genomes were explored to gain a global view of gene families among closely related species in Rosaceae. A total of 680,617 (94.4%) genes from BL1 and other 15 Rosaceae genomes were assigned to 46,182 gene families. There were 7,802 (16.89%) gene families shared by these genomes, implying their conservation. In the BL1 genome, 82,900 (94.23%) genes were assigned to orthogroups, and 2,019 orthogroups were species-specific, containing 6,188 genes. BL1 had the highest number of one-to-one orthologues with Hillquist and the lowest number of orthologues with flowering cherry (*Cerasus* x *kanzakura*) (Supplementary Table S4). Using the protein sequences of single-copy orthologs, a high-confidence phylogenetic tree was constructed (Figure 2). The diploid blackberry ‘Hillquist’ and tetraploid BL1 shared a common ancestor about 7.5 million years ago (MYA).

**Figure 2.**
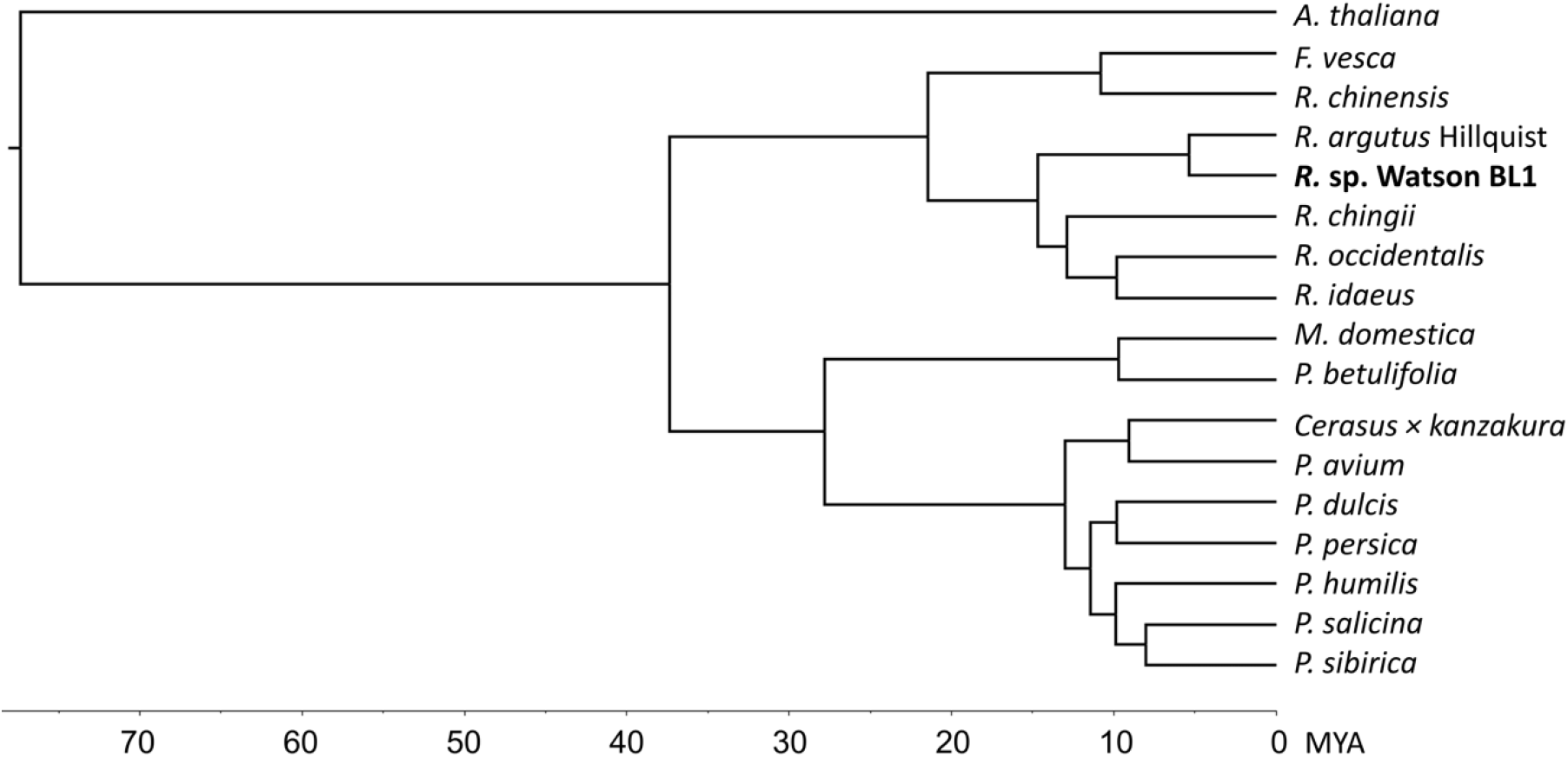
Phylogenetic tree of the BL1 blackberry and 15 plant species within the Rosaceae family inferred from published genome sequences. The genome of *Arabidopsis thaliana* was included as an outgroup. X-axis represents estimated divergence times (millions years ago, MYA) for the different lineages of Rosaceae plant species. *F*. *vesca* = *Fragaria vesca* (woodland strawberry, diploid) (Edger et al., 2018); *R*. *chinensis* = *Rosa chinensis* (‘Old Blush’, doubled haploid) (Hibrand Saint-Oyant, et al., 2018); *R*. *argutus* Hillquist = *Rubus argutus* ‘Hillquist’ (blackberry, diploid) (Brůna et al., 2023); *R*. sp. Watson BL1 = *Rubus* subgenus *Rubus* Watson BL1 selection (tetraploid); *R*. *chingii* = *Rubus chingii* (Fepenzi variety ‘Wanfu 1’, diploid) (Wang et al., 2021); *R*. *occidentalis* = *Rubus occidentalis* (black raspberry selection ORUS 4115-3, diploid) (VanBuren et al., 2016); *R*. *idaeus* (red raspberry variety ‘Anitra’, diploid) (Davik et al., 2022); *M*. *domestica* = *Malus domestica* (apple variety ‘Golden Delicious’, doubled haploid) (Daccord et al., 2017); *P*. *betulifolia* = *Pyrus betulifolia* (Oriental pear, diploid) (Dong et al., 2019); *Cerasus* × *kanzakura* (flowering cherry) (Shirasawa et al., 2021); *P*. *avium* = *Prunus avium* (sweet cherry, diploid) (?????); *P*. *dulcis* = (almond variety ‘Nonpareil’, diploid) (D’Amico-Willman et al., 2022) ; *P*. *persica* = *Prunus persica* (peach, diploid) (???); *P*. *humilis* = *Prunus humilis* (variety ‘Jing Ou No.2’, diploid) (Wang et al., 2022); *P*. *salicina* (Japanese plum variety ‘Sanyueli’, diploid) (Liu et al., 2020); and *P*. *sibirica* = *Prunus sibirica* (Siberian apricot) (Xu et al., 2022???).

A total of 2,019 orthogroups were unique to the BL1 genome, containing 6,188 genes. These unique genes in BL1 were further annotated using Blast2go. The highest number of direct gene ontology (GO) count unique to BL1 were found in biological categories such as cotyledon development, seed germination, epidermal cell differentiation, maintenance of floral organs, post-embryonic development, and regulation of cellular process (Supplementary Figure S10).

### 2.4 Chloroplast genome

The chloroplast genome of BL1 was assembled into a single contig of 156,514 bp with a GC content of 37.11% (Figure 3). The chloroplast genome contains 160 genes, including 97 protein-encoding genes, 10 rRNA genes, and 65 tRNA genes. The chloroplast genome displayed a typical quadripartite structure, with one large single copy (LSC) region (85,846 bp) and one small single copy (SSC) region (18,772 bp) separated by two inverted repeat (IR) regions (25,948 bp each). The BL1 chloroplast genome was in high concurrence with the published chloroplast genome of a hybrid blackberry ‘Arapahol’, which was 156,621 bp with 134 genes (Liu et al., 2021). For the different sequenced blackberry cultivars/lines in this research, the length of the plastomes ranged from 156,514 to 156,704 bp with similar GC contents (36.4%-37.4%). These chloroplast genome sizes are comparable to the chloroplast genomes of other *Rubus* sp. including *Rubus peltatus* (155,582 bp) (Qiao et al., 2022), *Rubus setchuenensis* (156,231 bp) (Zhu et al., 2022), and *Rubus lambertianus* (156,569 bp) (Chen et al., 2020).

**Figure 3.**
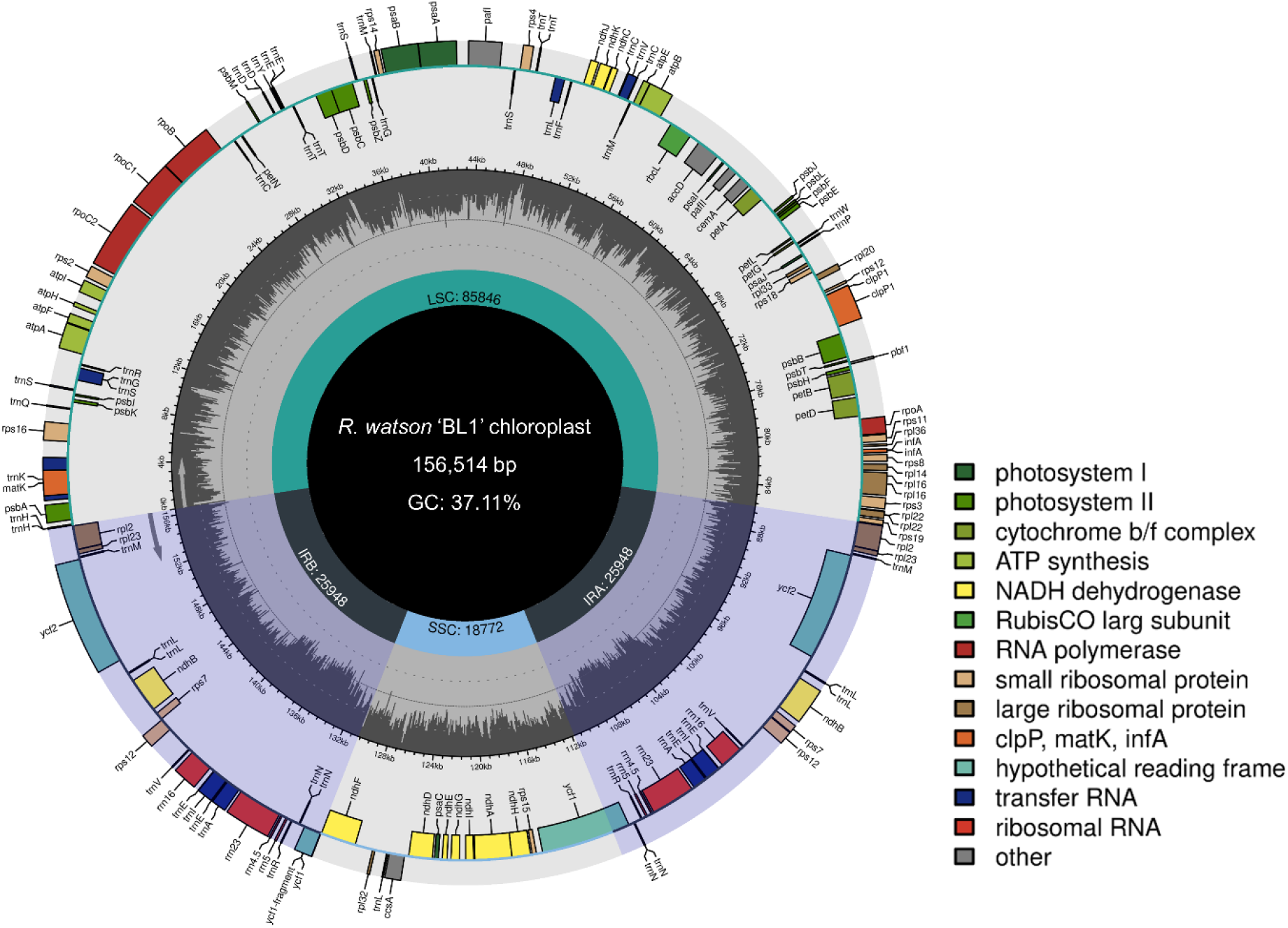
Chloroplast genome map of the BL1 blackberry with annotated genes. Genes inside the most outer circle are transcribed clockwise, and those outside the most outer circle are transcribed counterclockwise. Color coding of the genes is based on annotated functional groups. Boundaries of the small single copy (SSC) and large single copy (LSC) regions and inverted repeat (Ira and Irb) regions are denoted in the most inner circle.

### 2.5 Disease resistance and susceptibility genes

Blackberry cultivation, particularly in the subtropical regions, faces major challenges from disease pressure (Agehara et al., 2020; Marin et al., 2022). These diseases include leaf rust caused by *Kuehneola uredinis* (Marin et al., 2022), leaf spot caused by *Pseudocercospora pancratii* (Marin et al., 2023), Fusarium wilt caused by *Fusarium oxysporum* (Pastrana et al., 2020), downy mildew caused by *Peronospora sparsa* (Ali et al., 2019), powdery mildew caused by *Podosphaera aphanis* (Solano-Báez et al., 2022), and orange cane blotch caused by the algae *Cephaleuros virescens* (Browne et al., 2020). Developing disease-resistant varieties is crucial for helping growers address diseases in a socioeconomically and environmentally friendly manner. Therefore, identifying disease resistance genes is essential for facilitating breeding for disease resistance in blackberries (Afanador-Kafuri et al., 2015). The NLR class of *R* genes in plants are characterized by nucleotide-binding site (NBS) and leucine-rich repeat (LRR) domains, as well as a variable amino- and carboxy-terminal domains (DeYoung and Innes, 2006; McHale et al., 2006). Analyzing NLR genes is crucial as they represent the largest class of *R* genes and play a significant role in plant resistance to multiple diseases caused by bacterial, fungal, and viral pathogens. In addition, we investigated the *Mildew resistance Locus O* (*MLO*) genes, a type of disease susceptibility (*S*) genes whose mutation has been found to confer broad-spectrum resistance against powdery mildew (Kusch and Panstruga, 2017). Studying *MLO* genes is important because they offer an opportunity to edit *S* genes and develop varieties with improved resistance to this widespread disease.

A total of 856 transcripts (770 genes) were identified as NLR genes in the BL1 genome, with the highest number of NLR genes found in chromosome A2 (60, 7.7%), followed by B6 (57, 7.4%) (Figure 4). These NLR genes were further classified into different classes based on the presence of NB-ARC, Toll/Interleukin-1 receptor (TIR), coiled coil (CC), RPW8, and LRR domains (Figure 4). The largest class, with at least one NB-ARC domain and one of the other domains was NL (265, 34.41%), followed by TNL (160, 20.77%) (Supplementary Figure S11). The smallest class was NC, which contained the NB-ARC domain and the coiled coil (CC) domain (5, 0.64%). Most of these genes were clustered together (Figure 4), and all the chromosomes contained at least two NLR genes. Chromosomes A2 and B6 contained the highest number (60 and 57) of NLR genes, indicating that these chromosomes are crucial for disease resistance-related traits. The frequency of NLR domains was similar to that of the Hillquist genome, which contains a total of 228 genes with NLR domains, with NL and TNL as the largest classes (Supplementary Figure S12). The highest number of complete NLRs in the Hillquist genome was found in Ra02 (46, 20.17%), followed by Ra05 (40, 17.54%), Ra06 (33, 14.47%), and Ra07 (33, 14.47%) (Supplementary Figure S12).

**Figure 4.**
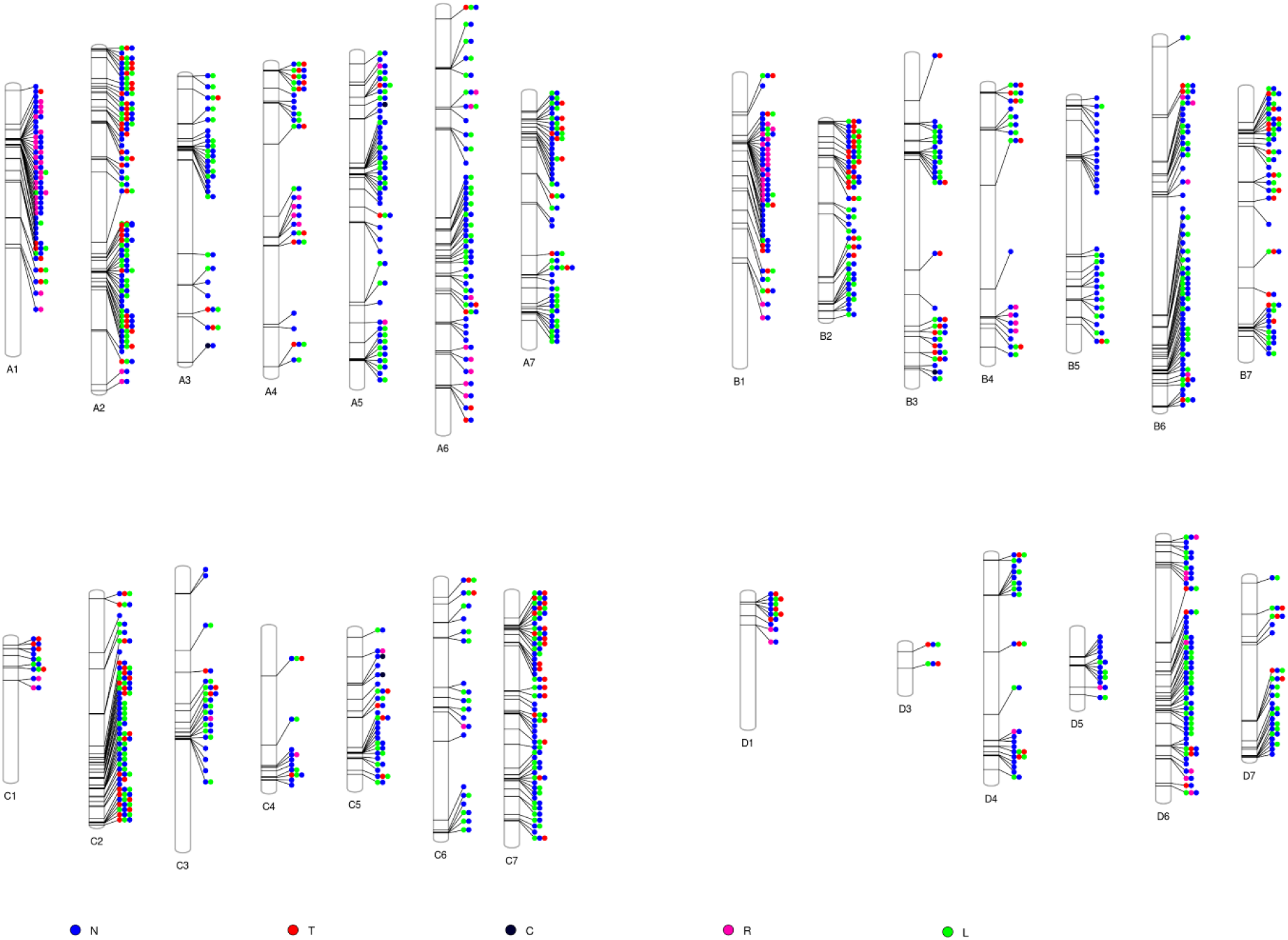
Distribution of NLR genes in the BL1 genome. Each vertical bar represents an assembled pseudochromosome in the BL1 genome. Blue dots represent genes with the NB-ARC domains, red dots the TIR domains, black dots the coiled coil (CC) domains, green dots the RPW8 domains, and lime dots the LRR domains.

A total of 74 genes were identified as members of the *MLO* family (Figure 1, Supplementary Figure S13). Twenty-one chromosomes each contain at least one *MLO* gene, with the highest number of *MLO* genes in Chromosome B1 and B5 (7 each). Homologs of all *AtMLO* genes, except *ATMLO7*, *ATMLO9*, *ATMLO10, ATMLO14*, and *ATMLO15*, were present in the BL1 genome, with *ATMLO6* having the highest number of homologous genes (19 in 11 chromosomes and 2 in contigs). *MLO* genes forms a relatively small family in blackberries compared to the NLR class of genes. Clade IV and clade V are known to be involved in susceptibility to powdery mildew in monocots and dicots, respectively (Kusch et al., 2016). In the BL1 genome, we did not identify any homologs of clade IV (Figure 5); however, 37 genes belonging to clade V were identified in the BL1 genome (Figure 5, Supplementary Table S5). These genes could act as susceptibility factors in blackberries and are potential candidates for follow-up studies to investigate if silencing, gene editing, or loss-of-function mutations in these genes could lead to powdery mildew resistance.

**Figure 5.**
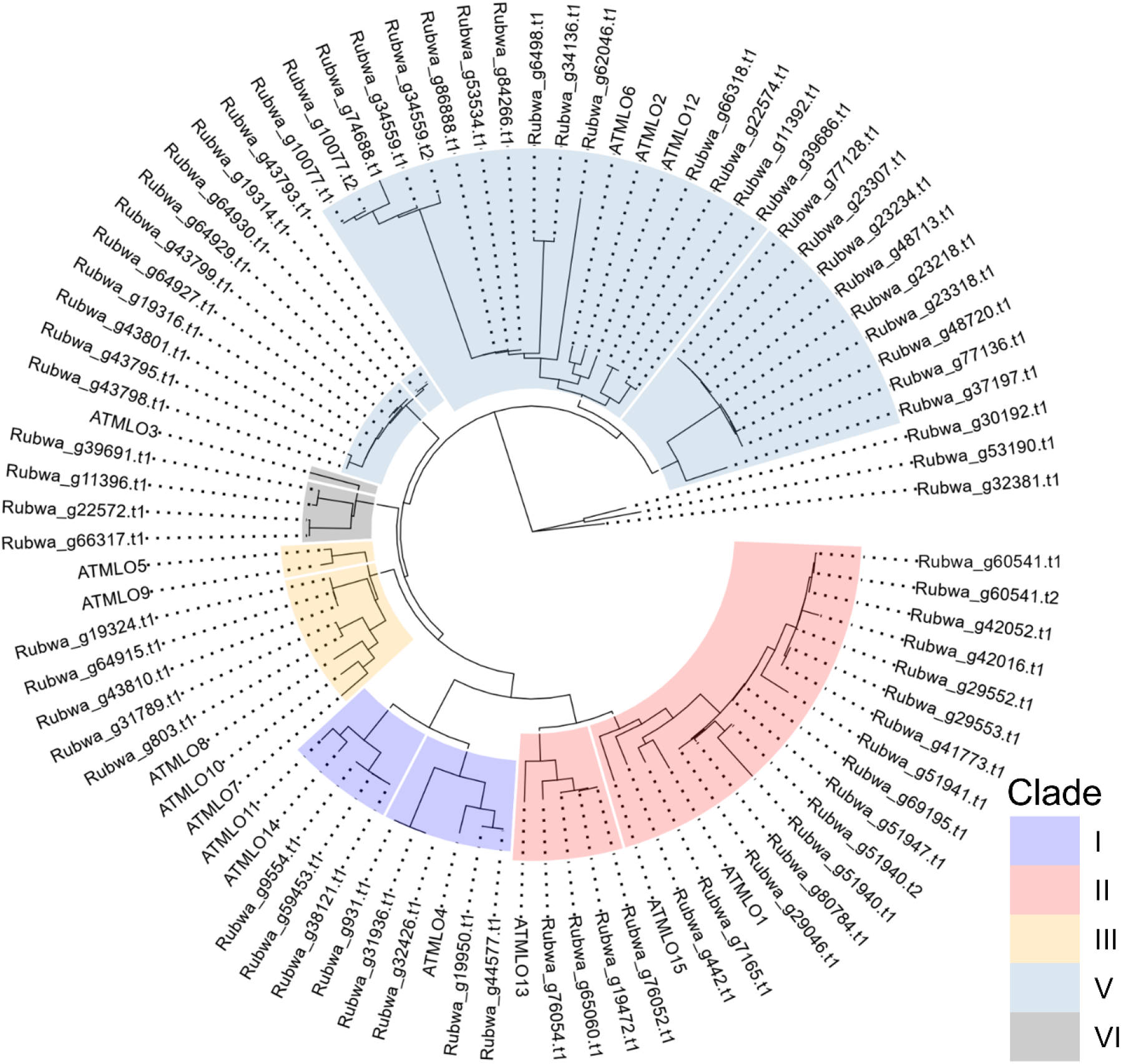
Clustering and phylogeny of *MLO* genes in the BL1 genome (prefixed with Rubwa) and *MLO* genes in *Arabidopsis thaliana* (prefixed with ATMLO).

### 2.6 Expression patterns of NLR and *DMR6* genes in blackberries

The BL1 genome assembly was utilized to understand the expression patterns of the NLR class of *R* genes and another *S* gene in different blackberry organs. We generated transcriptome sequencing data from thorny and thornless blackberry canes. In addition, blackberry transcriptome sequencing data were downloaded from four NCBI BioProjects (PRJNA680622, PRJNA701162, PRJNA744069, and PRJNA787794). In these BioProjects, transcriptome data were generated for blackberries in different developmental stages (green, red, and black; or unripe and ripe), or those treated with the plant growth regulator ABA, or blackberry leaf tissues collected from plants fertilized with different types of nitrogen fertilizers.

A total of 608 NLR genes exhibited expression in at least one of these RNA samples evaluated (Supplementary Figure S14, Supplementary Table S6). More NLR genes were expressed in blackberry leaves compared to fruit or shoot tissue. A higher number of genes were expressed in unripe berries compared to ripe berries. Similarly, more NLR genes were expressed in green and red berries as opposed to black berries.

A total of 4,046 transcripts, including 28 NLR genes, were upregulated in black berries, while 8,518 transcripts and 129 NLR genes were downregulated in black berries. In a separate experiment, 3,106 transcripts with 10 NLR genes were upregulated in ripe berries compared to 3,771 transcripts with 14 NLRs downregulated in ripe berries. A total of 1,333 transcripts were commonly upregulated in both black and ripe berries. Among them, two transcripts were from NLR genes (Supplementary Table S7). Similarly, 2,009 transcripts were commonly upregulated in both green and unripe berries, with 22 transcripts being NLR genes (Supplementary Table S7). The gene structure of these common transcripts revealed that most of the transcripts had a single exon (Supplementary Figure S15).

Blackberries are highly susceptible to downy mildew (DM) caused by *Peronospora sparsa* (Ali et al., 2019). The *Downy Mildew Resistance 6* (*DMR6*) gene is the *S* gene for DM in Arabidopsis (van Damme et al., 2008). This gene encodes 2OG-Fe(II) oxygenase (van Damme et al., 2008), and its mutation confers resistance to DM and other pathogens in Arabidopsis (van Damme et al., 2008; Zeilmaker et al., 2015). We mined the BL1 genome and identified 392 transcripts that contained the 2OG-Fe(II) oxygenase superfamily (PF03171) domain, out of which 350 also contained the DIOX_N domain (PF14226). A total of six *DMR6* transcripts were significantly upregulated in thorny blackberry shoots (Supplementary Table S8), while 26 *DMR6* transcripts were upregulated in black and ripe berries, and 14 *DMR6* transcripts were downregulated in black and ripe berries. The highest number of differentially expressed *DMR6* transcripts were present in Chromosome A2 (9 transcripts).

### 2.7 Candidate genes in the thorniness locus region

Most blackberry species exhibit sparse to abundant thorny protrusions on canes. These thorns, which are modified lateral branches, grow from lateral buds and are derived entirely from the tissues outside the vascular cortex (Clark et al., 2007). The crosstalk between auxin and cytokinin determines lateral bud activation, a precondition for thorn development (Müller and Leyser, 2011). Since domestication and commercial cultivation of blackberries, gardeners, horticulturalists, and growers have sought thornlessness to reduce injury risks during harvest and maintenance (Coyner et al., 2005). However, thornlessness has also associated with undesirable characters such as cold susceptibility, semi-erect growth habit, late-fruiting, and partial sterility (Hall et al., 1986a, 1986b).

A previous genetic mapping study identified one locus (*S* locus) for thorniness in blackberries (Castro et al., 2013). This locus was mapped to a region of 8.8 Mb in chromosome A4 from 30,682,793 to 39,471,371 bp of the BL1 genome (Figure 1). This genomic region contains 1,567 genes, with a majority involved in processes such as response to chemicals (6.42%), response to stress (6.04%), anatomical structure development (5.37%), biosynthetic process (4.62%), and multicellular organism development (4.43%) (Supplementary Figure S16). Among these genes, 49 genes have NLR domains, and two possess complete NLR domains (Rubwa_g15611.t1 and Rubwa_g15684.t1).

The thorniness locus region contains 255 transcripts with at least one HIGH impact variant (Supplementary Table S9). GO annotation of these transcript reveals a higher presence of those related to stress response, chemical response, anatomical structure development, biosynthetic process, and multicellular organism development (Supplementary Figure S17). Ninety-six of the transcripts within the locus region contain transcription factors (Supplementary Table S10), including bHLH and MYB, which are involved in trichome development (Pattanaik et al., 2014).

The bHLH family, a large group of transcription factors in *Arabidopsis*, include members such as *GLABROUS3* (*GL3*) and *ENHANCER OF GLABROUS3* (*EGL3*), which play a role in trichome development (Payne et al., 2000; Zhang et al., 2003). MYB16-like genes (MIXTA-like R2R3-MYB family member) are involved in trichome initiation and regulate conical cell outgrowth in diverse plant species (Oshima and Mitsuda, 2013). Chromosome 4 in red raspberry (*Rubus idaeus*) is also important for controlling prickle and shows differential expression of these transcription factors (Khadgi and Weber, 2021).

Several other important transcription factors, such as *AP2*, *DRF*, *MIKC*_*MADS*, *WRKY*, *NAC*, and *SBP*, were identified in this thorniness locus region of BL1 (Supplementary Figure S18). In particular, NAC transcription factors have been implicated in spine development in cucumbers (Liu et al., 2018), while SBP genes control initiation of trichomes in rice (Lan et al., 2019).

A comparative analysis of gene expression data revealed that 123 transcripts were upregulated in thornless blackberries, with four of these transcripts found within the thorniness locus region (Supplementary Table S11). Meanwhile, 520 transcripts were downregulated in thornless blackberries, and 14 of these transcripts were located within the thorniness locus region. These transcripts have been functionally characterized as being involved in trichome development in other species.

For instance, the pleiotropic drug resistance gene (*NpPDR1*) is constitutively expressed in the roots and leaf glandular trichomes of *Nicotiana plumbaginifolia*. These trichomes secrete terpenoid molecules as the first line of defense in response to plant pathogens (Stukkens et al., 2005). another differentially expressed gene, Caffeoyl-CoA, is involved in lignin biosynthesis in hops (Nagel et al., 2008).

A downregulated transcript, Glycosylphosphatidylinositol-anchored lipid protein transfer 1 (Rubwa_g15399.t1), has been found to be expressed in the trichomes in Arabidopsis leaves, and its disruption leads to alterations in cuticular lipid composition (Lee et al., 2009). In tomatoes, beta-ketoacyl reductase has been identified as trichome-enriched candidate gene (Ji et al., 2023).

The *DORNRÖSCHEN* (*DRN*) gene, also known as *ENHANCER OF SHOOT REGENERATION1* (*ESR1*) gene, contributes to the organization of meristems in Arabidopsis (Kirch et al., 2003) and plays a role in cytokinin-independent shoot regeneration (Banno et al., 2001). Interestingly, homocysteine S-methyltransferase 3, one of the top upregulated genes in the prickleless epidermis in *Solanum viarum* Dunal (Pandey et al., 2018), was found to be downregulated in thornless blackberries.

We identified four transcripts significantly upregulated in thornless blackberries, shedding lights on their potential roles in thorn development. Lysine histidine transporters are involved in importing amino acids into trichomes and play a role in organic N transfer for pollen production (Rabby et al., 2022). MYB domain protein 16 was also upregulated in thornless blackberries. The MIXTA-like transcription factor MYB16 is a major regulator of cuticle formation in vegetative organs (Oshima and Mitsuda, 2013). Additionally, the MED28 mediator subunit is essential for both development and senescence processes (Shaikhali et al., 2016). Lastly, one of the transcripts of Glycosylphosphatidylinositol-anchored lipid protein transfer 1 (Rubwa_g15399.t2) was found to be upregulated.

These up- and down-regulated genes are potential candidate genes for regulating the thornlessness trait in blackberries. These findings may hold considerable importance from a breeding perspective. Information for these genes is provided in Supplementary Table S10. In prickle-free epidermis tissue of red raspberry, transcription factors such as MIXTA-like R2R3-MYB family members, MADS-box, AP2/ERF, and NAC were significantly downregulated (Khadgi and Weber, 2021). We also identified these transcription factors in the thorniness locus region of blackberries (Supplementary Table S10, Supplementary Figure S18), suggesting that the mechanisms for thorn initiation in these two species could be similar. Furthermore, in the thornless blackberries, we observed downregulation of two other NLR transcripts (Rubwa_g6165.t in chromosome A2, and Rubwa_g54129.t1 in chromosome C1) and upregulation of two NLR transcripts (Rubwa_g33903.t1 in chromosome B1, and Rubwa_g79199.t1 in chromosome D6).

### 2.8 Candidate genes for primocane fruiting

The primocane fruiting trait was genetically mapped to one locus (*F* locus) between markers FF683693.1 Rh_MEa0007aG06 and FF683518.1 Rh_MEa0006aC04 in the blackberry genome map (Castro et al., 2013). This region corresponds to an 11 Mb segment in *R. argutus* chromosome Ra02 at 25,901,374 to 25,901,083 bp (FF683518.1 Rh_MEa0006aC04) and 37,085,586 to 37,085,204 bp (FF683693.1 Rh_MEa0007aG06) (Brůna et al., 2023). In the BL1 tetraploid genome, Ra02 corresponds to Chromosome A2, and the region could be mapped to a 12.5 Mb segment between 29,653,983 and 42,218,375 (Figure 1). This region contains 2,172 genes.

Comparison with the FLOR-ID database (Bouché et al., 2016) revealed 32 flowering-related homologs within this locus region (Supplementary Table S13). These genes have been reported to have positive effects on flowering time, such as *FT*-*INTERACTING PROTEIN 1, IDD8*, *GA1*, *GID1B*, *LD*, *NF*-*YB2*, *SUS4*, *SKB1*, *FPF1*, *DCL3*, *MRG1*, and *GA20OX3*. Additionally, genes with negative effects on flowering time, such as *HTA*, *WDR5A*, *CSP2*, *TPL*, *UBP26*, and *GASA5*, were present in the locus region. A loss-of-function mutation in a repressor could result in recessive inheritance of this trait, making floral repressors the primary candidates for primocane fruiting (Brůna et al., 2023). As a result, these identified genes might play a critical role in primocane fruiting in blackberries and merit further investigation. We identified four genes encoding transcription factors such as C2H2 (Rubwa_g6736.t1), HB-other (Rubwa_g7150.t1), and NF-YB (Rubwa_g7154.t1) in the primocane-fruiting locus. Additionally, genes involved in the photoperiod pathway, including *FTIP1* (Rubwa_g6515.t1, Rubwa_g6515.t2, Rubwa_g6518.t1), *NF-YB* (Rubwa_g7154.t1), and *TPL* (Rubwa_g7556.t1, Rubwa_g7559.t1), were also identified within this locus region.

### 2.9 Sequence polymorphisms among tetraploid blackberry cultivars/selections

We resequenced the genomes of seven other blackberry cultivars/selections (‘Kiowa’, ‘Osage’, ‘PrimeArk® 45’, ‘PrimeArk® 45 variant with fewer and shorter thorns, ‘PrimeArk® Freedom’, ‘PrimeArk® Traveler’, and BL2 selection) to uncover sequence polymorphisms that may contribute to their diverse characteristics. In our analysis, we identified a total of 13,378,961 genetic variants, with an average rate of one variant every 65 bp (Supplementary Table S14). The majority of these variants were classified as SNPs (9,846,835, 73.59%), with insertions and deletions accounting for 9% (573,329 insertions and 641,516 deletions). Most SNP effects were classified as “MODIFIER” putative impact (97.914%), while 36,553 SNP (0.072%) effects were considered “HIGH” putative impact, 464,161 SNP (0.909%) effects as “LOW” impact, and 564,543 (1.106%) effects as “MODERATE” putative impact. SnpEff also classified 463,095 (53.00%) as missense, 8,004 (0.92%) as nonsense, and 402,724 (46.09%) effects as silent functional classes. Most of the variants were located in intergenic regions (21.428%) or upstream and downstream regions (32.191% and 31.452%, respectively), with 1.97% of the variants found in exons. Based on nucleotide substitutions, 20,524,771 SNPs were classified as transitions, and 11,448,387 SNPs were classified as transversions, resulting in a genome-wide transition to transversion ratio (Ts/Tv) of 1.79. A summary of the variants for individual blackberry samples is provided in Supplementary Table S14.

By comparing the variant information of leaf rust-susceptible cultivars ‘Kiowa’ and ‘Prim-Ark® Freedom’ to leaf rust-resistant cultivars ‘Osage’, ‘Prim-Ark® Traveler’, and ‘Prim-Ark® 45’, we identified 284 variants present in 54 NLR genes (Supplementary Table S15) that exhibited differences between leaf rust-resistant and susceptible cultivars. Notably, two SNPs were found in Rubwa_g15684.t1, located in the thorniness locus region. The analysis of these genetic differences may be crucial for understanding the mechanisms underlying disease resistance and thornlessness.

## 3 Conclusion

In this study, we present the first chromosome-length, phased genome assembly and annotation of the tetraploid blackberry selection BL1. Comparisons with the published diploid Hillquist genome attest to the high quality of the BL1 genome, while BUSCO analysis further demonstrates its high completeness and comprehensiveness. Using comparative genomic approaches, we shed light on the relationship between BL1 with other 15 sequenced genomes within the Rosaceae family and revealed that diploid and tetraploid blackberries might diverged at approximately 7.5 MYA. We identified numerous disease resistance and susceptibility genes (NLR, *MLO*, and *DMR6*) in the BL1 genome, which will be instrumental in developing disease-resistant blackberry cultivars. Moreover, we discovered candidate genes and transcription factors associated with thornlessness, which will aid in the development of thornless blackberry cultivars. Resequencing seven blackberry cultivars/selections unveiled genetic diversity that can be harnessed for future breeding efforts. The BL1 genome sequence and annotation will serve as an invaluable resource for breeding, genetic, genomic, and molecular research in blackberries and related berry crops. The release of this tetraploid blackberry genome can contribute to more efficient and targeted breeding, ultimately leading to the development of new cultivars with enhanced fruit quality, desirable fruiting habits, as well as resistance to important diseases.

## 4 Materials and methods

### 4.1 Plant material and experimental design

The BL1 blackberry selection was chosen for genome sequencing and assembly because it resulted from a cross between the first two important commercial thornless, primocane-fruiting cultivars ‘Prime-Ark® Traveler’ (PAT) (Clark and Salgado, 2016) and ‘Prime-Ark® Freedom’ (PAF) (Clark, 2014), and exhibited multiple desirable traits, including thornless canes, primocane-fruiting, and good adaptation to subtropical growing conditions. ‘Prime-Ark® Freedom’ was the world’s first commercially released thornless primocane-fruiting blackberry, while ‘Prime-Ark® Traveler’ is the first thornless primocane-fruiting blackberry cultivar with good shipping quality. Both PAT and PAF were included in the genome resequencing. In addition, three other blackberry cultivars and two selections were included for genome re-sequencing, and they were ‘Kiowa’ (Moore and Clark, 1996), ‘Osage’ (Clark, 2013), ‘Prime-Ark® 45 (PA45) (Clark and Perkins-Veazie, 2011), one selection of PA45 with fewer and shorter thorns (S. Parajuli and Z. Deng, unpublished), and the BL2 selection. These cultivars/selections represent substantial variations in thornlessness, fruiting habit, and resistance to leaf rust disease (Supplementary Table S16).

### 4.2 Estimation of genome size

Young unexpanded leaves of BL1 blackberry were used for flow cytometry. The one step protocol with Tris MgCl_2_ lysis buffer, RNase, and propidium iodide recommended by Doležel et al. (2007) was followed for sample preparation. ‘Polanka’ soybean (*Glycine max* Merr.; 2.50 pg/2C) and ‘Stupické polní rané’ tomato (*Solanum lycopersicum* L.; 1.96 pg/2C) were used as internal standards, and three technical replicates were analyzed for each of the three biological replicate samples. Holoploid DNA content (2C) was calculated as DNA content of standard x mean fluorescence value of sample / mean fluorescence of internal standard. Monoploid genome sizes were calculated by dividing the 2C genome size by the inferred ploidy (Rothleutner et al., 2016).

Chromosome squashing was performed on dividing root tips from BL1 plants. Root tips (approximately 1 cm) were isolated and immersed in 2 mM 8-hydroxyquinoline for 3 hours at 4 °C. Pre-treated roots were washed with deionized water and placed in a fixative solution (3 methanol:1 glacial acetic acid, v/v) at 4 °C overnight. The next day, roots were washed again in deionized water and placed in 1N HCl for 22 min. Acid was removed and root tips were washed a final time in deionized water and stained in 2% acetocarmine solution (Carolina Biology Supply Company) overnight. Root caps and non-meristematic tissue were removed under a stereoscope. Root tips were squashed in a drop of acetocarmine solution with a cover slip on a glass slide. Slides were viewed under a bright field microscope at ×1000 magnification.

### 4.3 DNA extraction and genome sequencing

Young leaf tissues were collected from BL1 and several other blackberry genotypes and immediately put into liquid nitrogen. Frozen leaf tissue samples were shipped on dry ice to NextOmics Biosciences (Wuhan, China) where DNA isolation and sequencing were performed. High-molecular weight DNA was extracted using the Qiagen DNeasy® Plant kit. For ONT sequencing, sequencing libraries were prepared following the Oxford Nanopore Technologies’ (Oxford Science Park, Oxford, UK) 1D genomic DNA by ligation (SQK-LSK109) – PromethION (version GDE_9063_v109_revD_04Jun2018) protocol and sequenced on the Nanopore PromethION platform. For Illumina sequencing, the Illumina TruSeq DNA PCR-free LPP (revision A, January 2013, low sample with 550 bp insert size) protocol was followed, and PCR-free sequencing libraries were sequenced on a NovaSeq 4000.

### 4.4 Sequence data processing and genome assembly

The quality of ONT data was assessed using NanoPlot/1.30.1. Raw reads were trimmed to a –min_tirm_size of 6 using porechop/0.2.4. Several assemblers were evaluated for assembling these nanopore sequencing data, including canu/1.8, wtdbg/2.5, WENGAN/0.2, SMARTdenovo, NECAT, and Flye/2.91.1. A summary of the assemblies is provided in Supplementary Table S17. Parameter optimization was performed for NECAT, and the assembly with the longest total length (setting3) and highest N50 (default) were chosen for Hi-C scaffolding (Supplementary Table S18). The final assembly was produced by Hi-C scaffolding on contigs of NEC–T - setting3.

### 4.5 Genome anchoring and scaffolding

Chromatin conformation capture data were generated using the Phase Genomics (Seattle, WA) Proximo Hi-C 4.0 Kit, which is a commercially available version of the Hi-C protocol. Following the manufacturer’s instructions for the kit, intact cells from blackberry leaf tissues were crosslinked using a formaldehyde solution, digested using a cocktail of restriction enzymes (*DPN*II, *Dde*I, *Hin*fI, and *Mse*I), end repaired with biotinylated nucleotides, and proximity ligated to create chimeric molecules composed of fragments from different regions of the genome that were physically proximal in vivo, but not necessarily genomically proximal. Continuing with the manufacturer’s protocol, DNA molecules were pulled down with streptavidin beads and processed into an Illumina-compatible sequencing library. Sequencing was performed on an Illumina NovaSeq sequencer, generating a total of 78,642,356 PE150 read pairs.

Reads were aligned to the BL1 draft genome assembly from above, also following the manufacturer’s recommendations. Briefly, reads were aligned using BWA-MEM with the -5SP and -t 8 options specified, and all other options default. SAMBLASTER was used to flag PCR duplicates, which were later excluded from analysis. Alignments were then filtered with samtools using the -F 2304 filtering flag to remove non-primary and secondary alignments. Putative misjoined contigs were broken using Juicebox based on the Hi-C alignments. A total of 101 breaks in 90 contigs were introduced, and the same alignment procedure was repeated from the beginning on the resulting corrected assembly.

The Phase Genomics’ Proximo™ Hi-C genome scaffolding platform was used to create chromosome-scale scaffolds from the corrected assembly, as described in Bickhart et al. (2017) As in the LACHESIS method, this process computed a contact frequency matrix from the aligned Hi-C read pairs, normalized by the number of restriction sites on each contig, and constructed scaffolds in such a way as to optimize expected contact frequencies and other statistical patterns in Hi-C data. Approximately 60,000 separate Proximo runs were performed to optimize the number of scaffolds and scaffold construction in order to make the scaffolds as concordant with the observed Hi-C data as possible.

### 4.6 Annotation of repetitive elements

For the annotation of repetitive elements (TEs), RepeatModeler/2.0 was used to create a *de novo* TE library. Miniature inverted transposable elements (MITEs) were identified using MITE-Hunter. LTR retrotransposons were detected using LTR_finder and LTR_harvest. LTR_retriever was utilized to process the outputs, and the predicted TEs were integrated into RepBase. RepeatMasker/4.1.1 was then used for TE annotations. The Telomere Identification toolkit (tidk) (https://github.com/tolkit/telomeric-identifier) was employed to identify telomeric repeat sequences, searching for the canonical repeat unit for the Rosales clade: AAACCCT.

### 4.7 Gene prediction and annotation

Gene prediction was performed using the BRAKER/v2 pipeline (Hoff et al., 2016), utilizing the transcriptome generated from RNA-Seq of thorny shoots, thornless shoots, leaves, green fruits, and publicly available RNA-Seq datasets from PRJNA680622, PRJNA701162, PRJNA744069, and PRJNA787794. BRAKER was run with protein data from OrthoDB for Viridiplantae. The final gene predictions, which combined gene prediction supported by both RNA-Seq and homologous protein evidence, were extracted using TSEBRA that (Gabriel et al., 2021).

### 4.8 Chloroplast genome assembly and annotation

The chloroplast genome of BL1 was assembled using GetOrganelle v 1.7.5.0 (Jin et al., 2020) with Illumina short reads. GeSeq (Tillich et al., 2017) was employed for annotating the chloroplast genome, and the resulting plot was generated using Chloroplot (Zheng et al., 2020).

### 4.9 RNA isolation, RNA-Seq library preparation and sequencing

RNA extraction, mRNA enrichment, RNA-Seq library construction, and sequencing were performed by NextOmincs Biosciences and CD Genomics (Shirley, NY, USA). RNA was extracted from each samples using the RNeasy kit (Qiagen, Switzerland) following the manufacturer’s protocol. Two micrograms of RNA from each sample were used to prepare RNA-Seq libraries, and then the libraries were sequenced on the Illumina NovaSeq4000 (NextOmics) or NextSeq (CD Genomics). Publicly available RNA-Seq datasets were downloaded from PRJNA680622, PRJNA701162, PRJNA744069, and PRJNA787794.

### 4.10 RNA-Seq expression analysis

Differential expression analysis was done by comparing thorny and thornless shoots of blackberry line PA45m. A False Discovery Rate of 0.05 was used, and the transcripts were filtered for a log2 fold change of ±2.

### 4.11 Analysis of genome resequencing data

Illumina sequences from seven other blackberry genotypes were processed using trimmomatic (Bolger et al., 2014) to remove adapters and low quality reads. The processed reads were then aligned to the BL1 assembly using HISAT2. Freebayes (Garrison and Marth, 2012) was employed to call SNPs. The vcf file was filtered to maintain a minimum depth of 15 and minimum quality of 20. All filtered SNPs were input into SnpEff (Cingolani et al., 2012) to predict the effects of SNPs.

### 4.12 Comparative genomics

The predicted proteins from the BL1 blackberry and other related Rosaceae genomes [woodland strawberry: *Fragaria vesca*; diploid blackberry: *Rubus argutus* cv. Hillquist; red raspberry: *Rubus idaeus* cv. Anitra; rose: *Rosa chinensis* ‘Old Blush’; *R*. *chingii*; and black raspberry: *R. occidentalis*) were grouped into orthogroups using OrthoFinder/2.5.2. A high confidence phylogenetic tree was constructed using the protein sequences of 1,847 single-copy orthologs with OrthoFinder/2.5.2. To calibrate the tree, we used r8s (Sanderson, 2003) and fixed the divergence time between *Rosa’* and *Fragaria* at 27.1 MYA, as suggested by the TimeTree website (https://timetree.org).

### 4.13 Identifying disease resistance and susceptibility genes

For identification of disease resistance genes, we scanned the predicted blackberry protein domains using the NBS Hidden Markov Model (HMM) profile with hmmer/3.2.1. Protein domains were predicted using the following Pfam HMMs: NB (Pfam accession PF00931), TIR (PF01582), RPW8 (PF05659), and LRR (PF00560, PF07725, PF13306, PF13855) domains, which were downloaded from the Pfam website (http://pfam.xfam.org/). Pfam hits were filtered using P <1e-04. To identify potential coiled-coil (CC) structures, we used DeepCoil and set the threshold for CC structure detection at 0.82. For phylogenetic tree construction of NBS-encoding genes, we first aligned the proteins with complete NBS domains using MAFFT. We then utilized the alignment for tree construction with RAxML and visualized the tree using MEGA7. We scanned blackberry protein domains for the MLO HMM profile using hmmer/3.2.1 for domain PF03094.

### 4.14 Transcription factor prediction

Transcription factors were identified using the Plant Transcription Factor Database (PlantRegMap).

## 5 Acknowledgements

We thank the following sequencing companies for genome and transcriptome sequencing or contig scaffolding: NextOmics Biosciences (Wuhan, China) for Oxford Nanopore Sequencing, Illumina sequencing, and RNA-Seq, CD Genomics (Shirley, NY, U.S.A.) for Illumina sequencing and RNA-Seq, DoveTail Genomics (Sotts Valley, CA, U.S.A.) for Hi-C analysis, and PhaseGenomics (Seattle, WA, U.S.A.) for Hi-C and scaffolding. All genome and transcriptome assembly and analysis were conducted on the University of Florida (UF) supercomputer (HiPerGator) that was provided by the UFIT Research Computing unit (Gainesville, FL, U.S.A.). We are grateful to Dr. Jaroslav Doležel (Institute Experimental Botany, Olomouc, Czeck Republic) for providing ‘Polanka’ soybean (*Glycine max* Merr.) and ‘Stupické polní rané’ tomato (*Solanum lycopersicum* L.) that were used in flow cytometrical analysis as internal standards.

## 6 Contributions

SP collected blackberry leaf tissue samples, isolated genomic DNA for genome sequencing or resequencing, isolated RNA for transcriptome sequencing, and prepared blackberry leaf tissues for Hi-C genome scaffolding; ZP coordinated genome and transcriptome sequencing with sequencing companies; DP assembled, annotated, and analyzed the blackberry genome and transcriptomes, identified candidate genes, conducted phylogeny analysis, and wrote the manuscript; SBP performed chromosome squashing, ran qRT-PCR, and analyzed gene expression; ZD initiated and supervised the project, and revised the manuscript. All authors reviewed the manuscript.

## 7 Data availability statement

The assembly and annotation files of the BL1 blackberry genome are uploaded to the Rosaceae database (xxxx).

