## Supplementary Figure for "A chromosome-scale and haplotype-resolved genome assembly of tetraploid blackberry (*Rubus* L. subgenus *Rubus* Watson)"

### Supplementary Figures

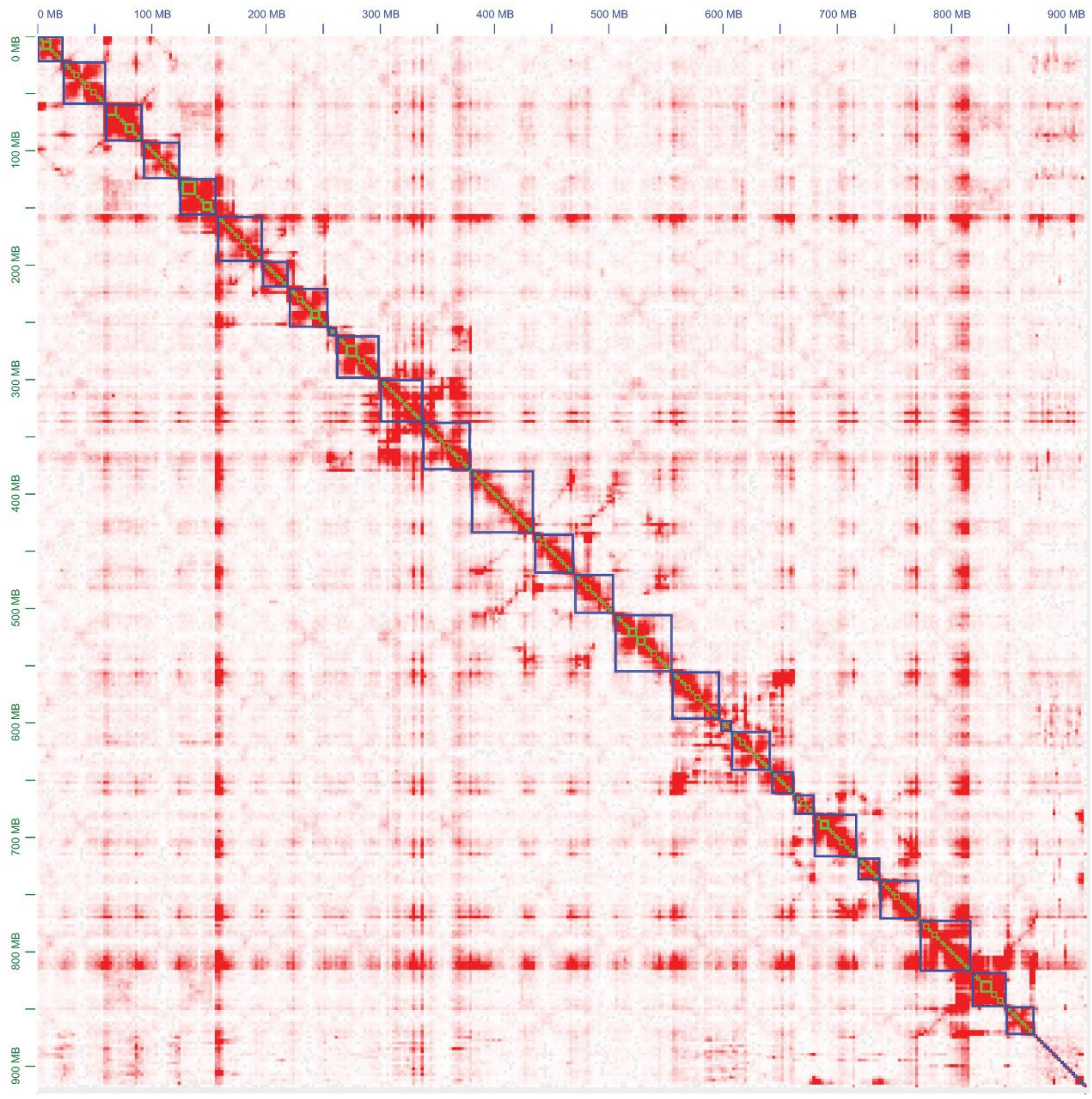


**Supplementary Figure S1.** Genome-wide Hi-C contact map of the hybrid BL1 (*Rubus* L. subgenus *Rubus* Watson) v1 assembly. The X and Y axes represent the coordinates of the chromosomes and the color scale reflects the frequency of contacts between two regions of the genome (arbitrary units), from white (rare contacts) to dark red (frequent contacts) at the 2.5-Mb matrix resolution. The 27 chromosomes and 328 unplaced scaffolds are numbered and oriented according to the diploid blackberry Hillquist genome in copies of four for each sub-genome and are presented as blue boxes in descending order (from upper left to lower right). Contacts (red pixels) within chromosomes are denser than between chromosomes, with contacts between adjacent genomic loci (pixels nearer to the diagonal) being denser than those at greater inter-locus distances (pixels farther off the diagonal). This feature of Hi-C was exploited to perform the scaffolding.

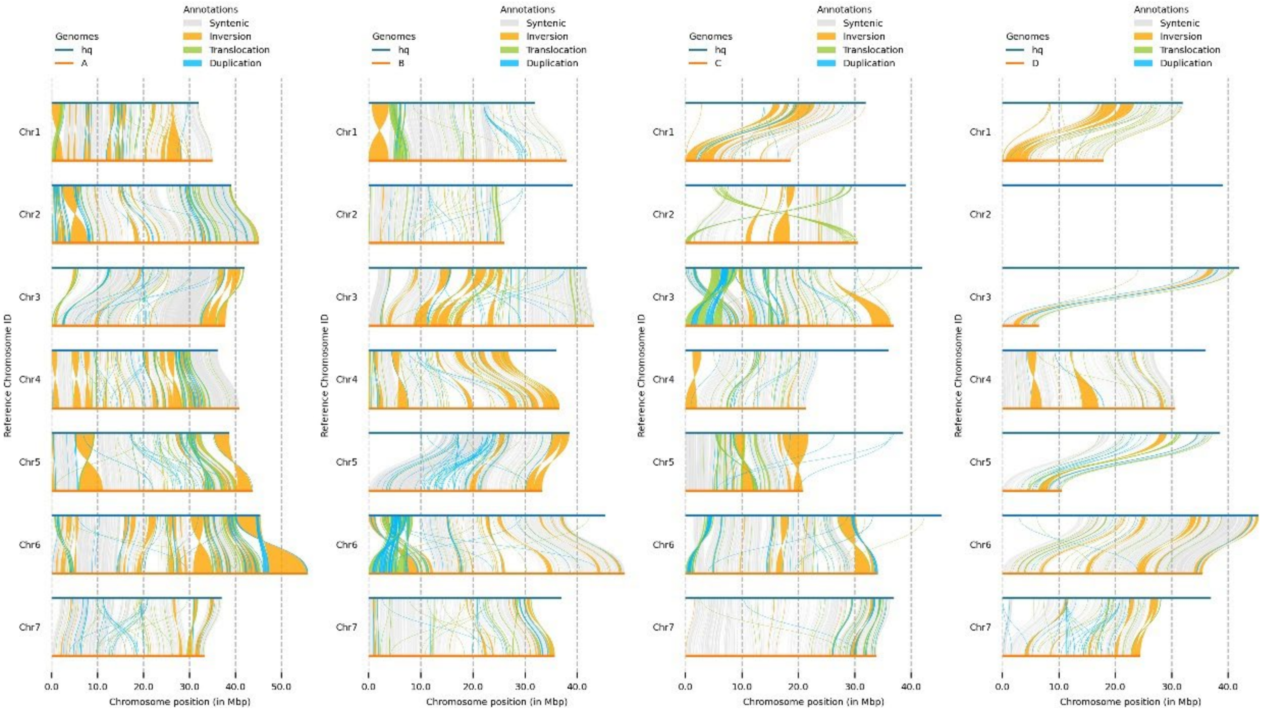


**Supplementary Figure S2.** Collinearity between the chromosomes of the Hillquist genome (hq: horizontal blue bars) and the four sub-genomes (A, B, C, and D) of the BL1 genome (horizonal orange bars). Chromosomal arrangements are shown with different colors: syntenic (platinum), inversion (saffron), translocation (pistachio), and duplication (blue).

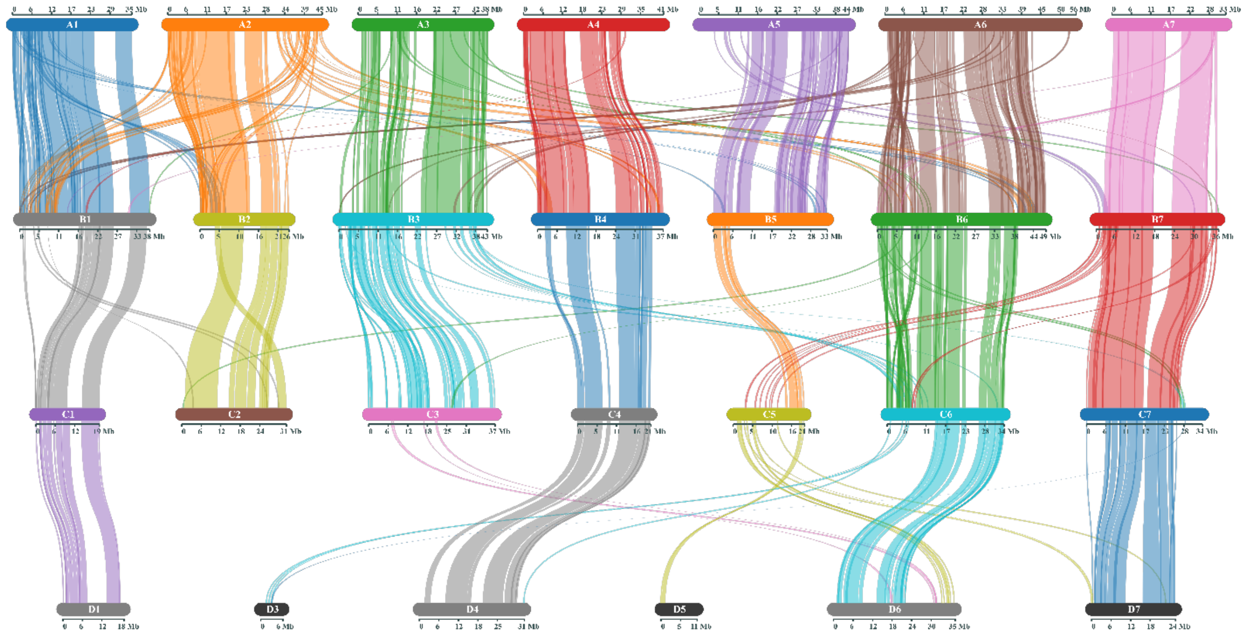


**Supplementary Figure S3.** Collinearity among the homoeologous chromosomes in the assembled tetraploid blackberry BL1 genome. The lines represent conserved gene arrays between chromosomes. Chromosomes were drawn proportionally with respect to the number of genes on each chromosome.

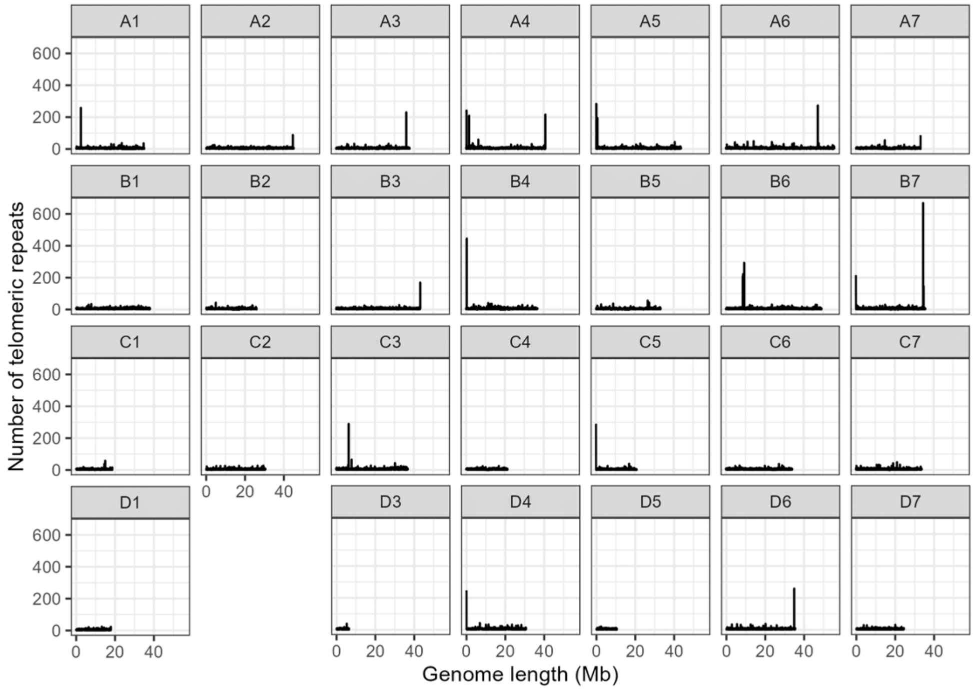


**Supplementary Figure S4.** Telomeric repeats identified in the chromosomes of the BL1 blackberry. The x-axis represents genome length and y-axis represents number of telomeric repeats identified. Higher number of repeats are present in the telomeric region.

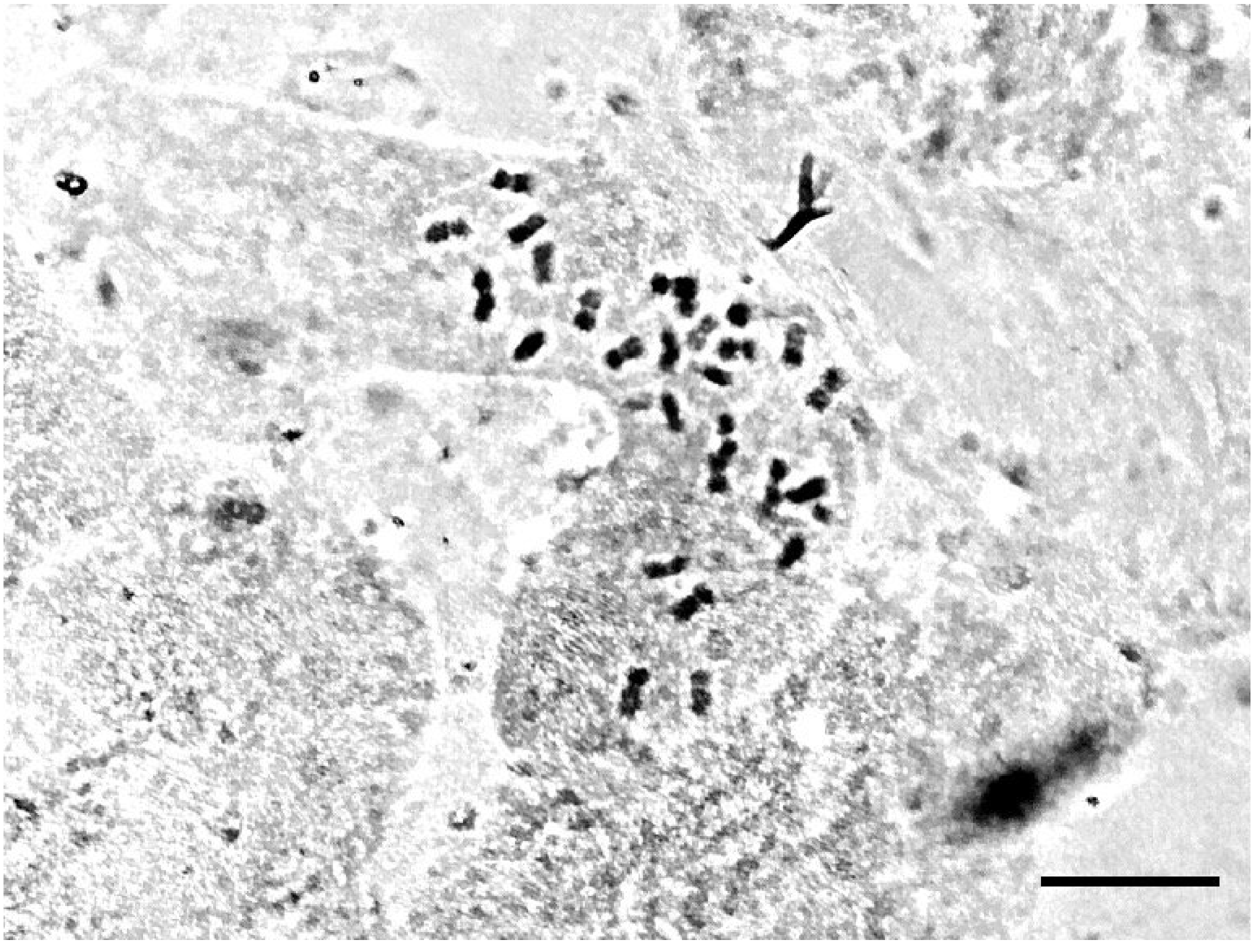

**Supplementary Figure S5**. Metaphase chromosomes ($2n = 4x = 28$) of the BL1 blackberry stained in acetocarmine solution. Scale bar = 2 µm.

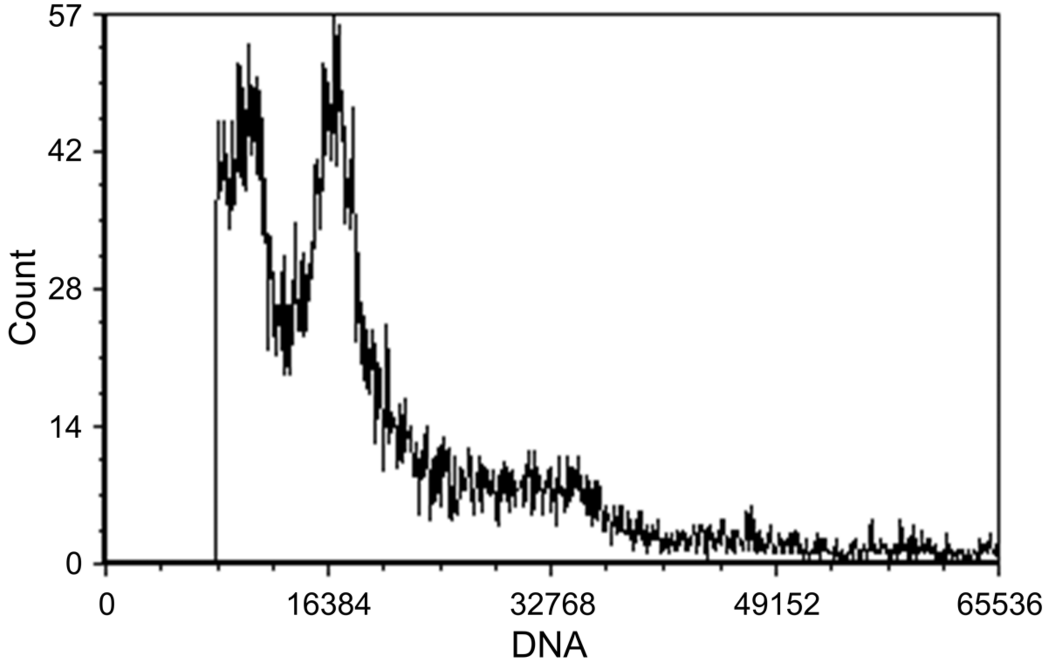


**Supplementary Figure S6.** Estimation of nuclear DNA content of the BL1 blackberry using flow cytometry. The left peak contains nuclei of the BL1 blackberry (1.46 pg/2C) and the right peak contains nuclei of 'Polanka' soybean (2.50 pg/2C).

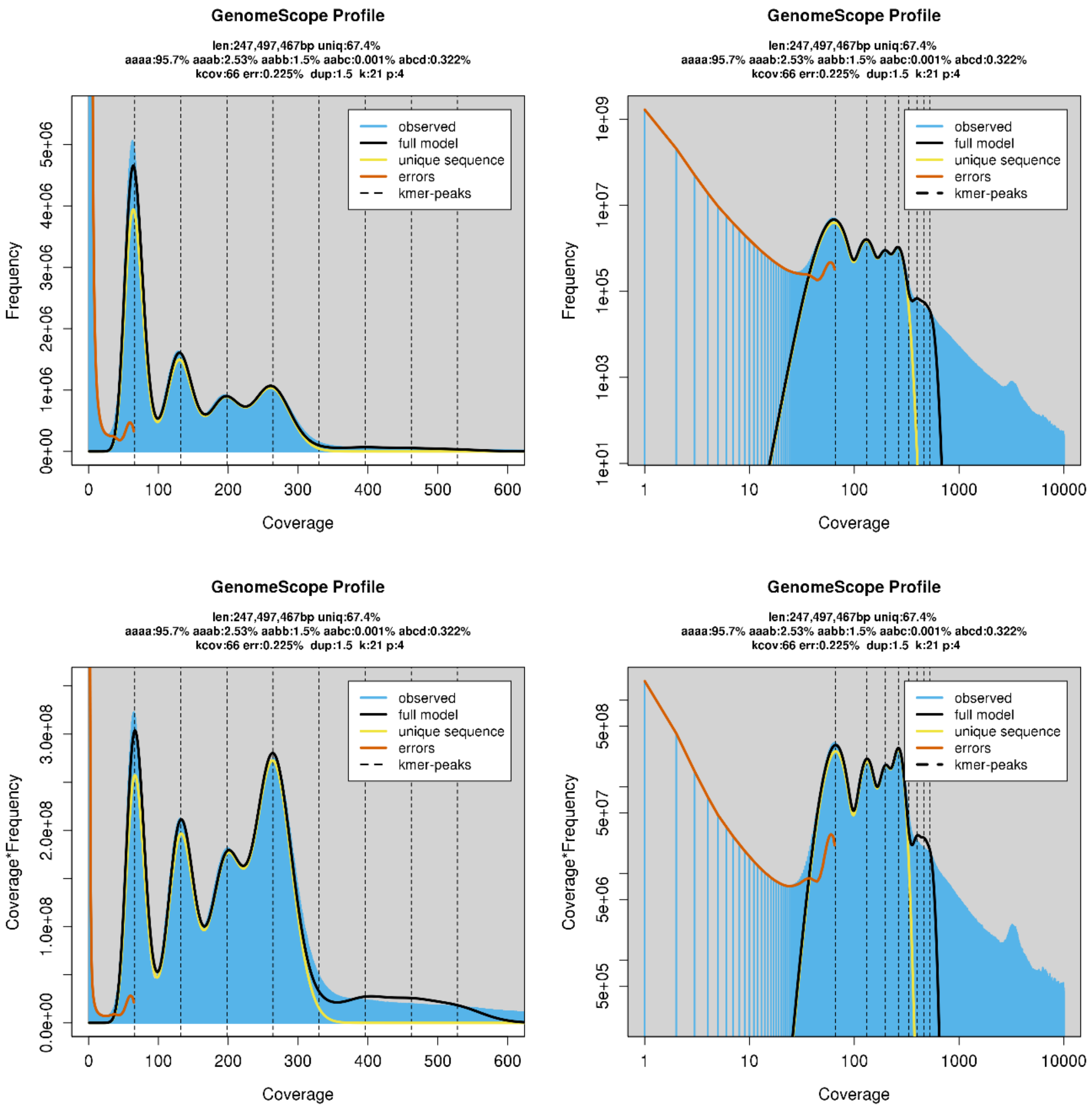


**Supplementary Figure S7.** GenomeScope results. Plots of the best fit model overlaying the *k*-mer spectrum for $k = 21$ for (A) untransformed linear (top left), (B) untransformed Log (top right), (C) transformed linear (bottom left), and (D) transformed Log (bottom right). The autotetraploid plot shows *aaab* > *aabb*.

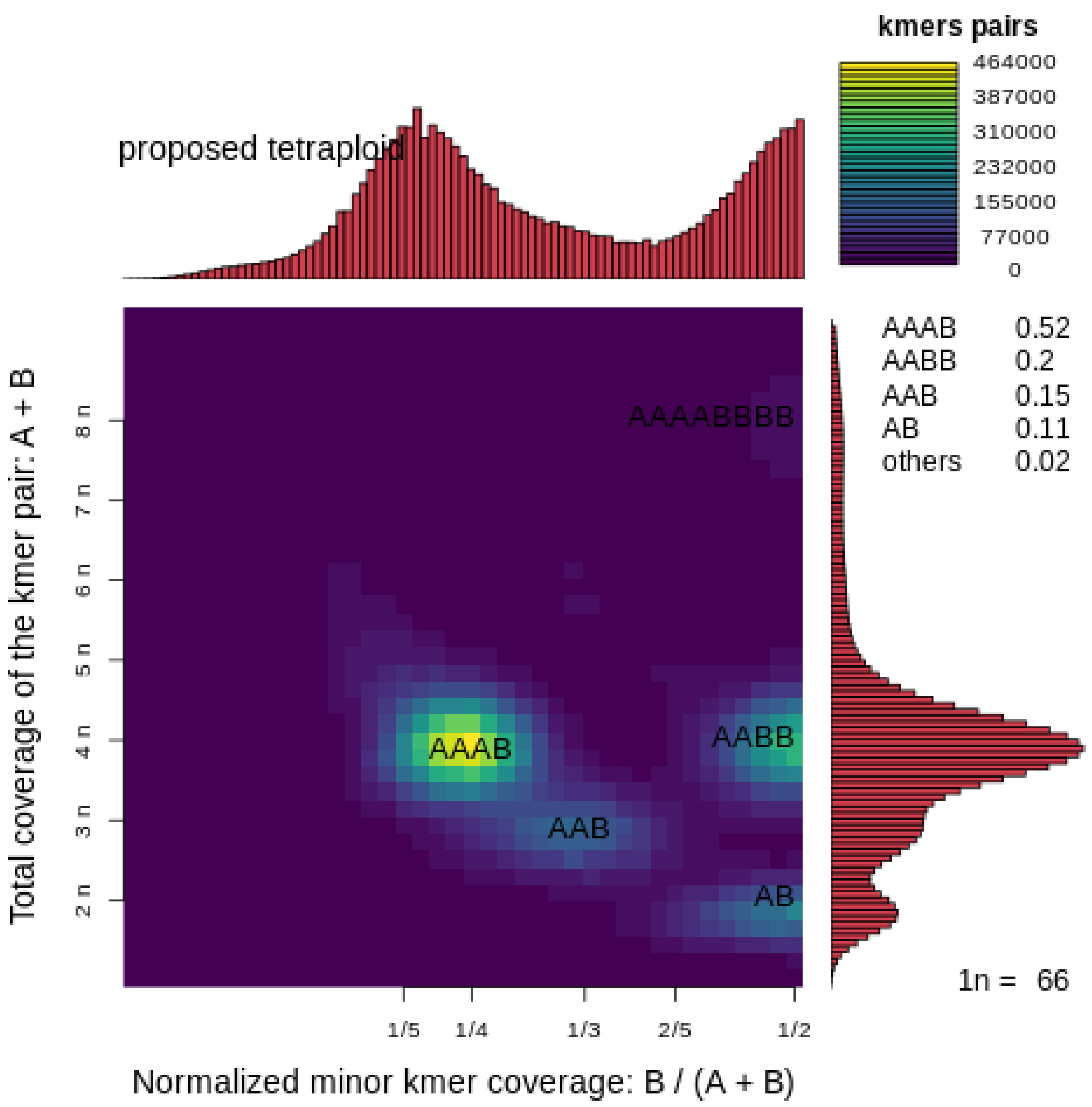


**Supplementary Figure S8.** Smudgeplot for the BL1 genome. Jellyfish determined k-mer pairs were used to determine the brightness of each smudge with the coloration indicating the approximate number of k-mer pairs per bin. The brightest smudge 'AAAB' (0.52) suggests that BL1 is an auto-tetraploid.

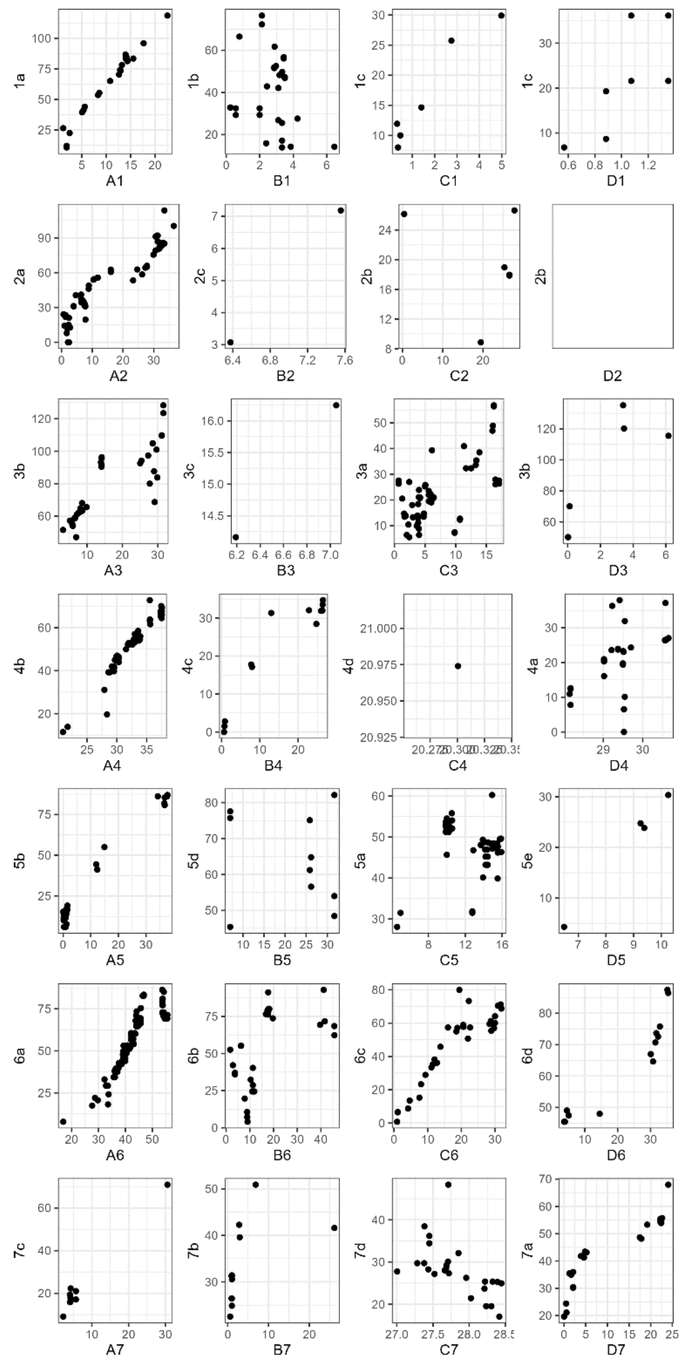


**Supplementary Figure S9.** Dotplot showing chromosomes of the BL1 genome on the x-axis and the blackberry linkage map (Brůna et al., 2022) on the y-axis. Sequences flanking the SNPs on the blackberry linkage map were mapped to the BL1 genome, and the dots represent 100% match of these sequences.

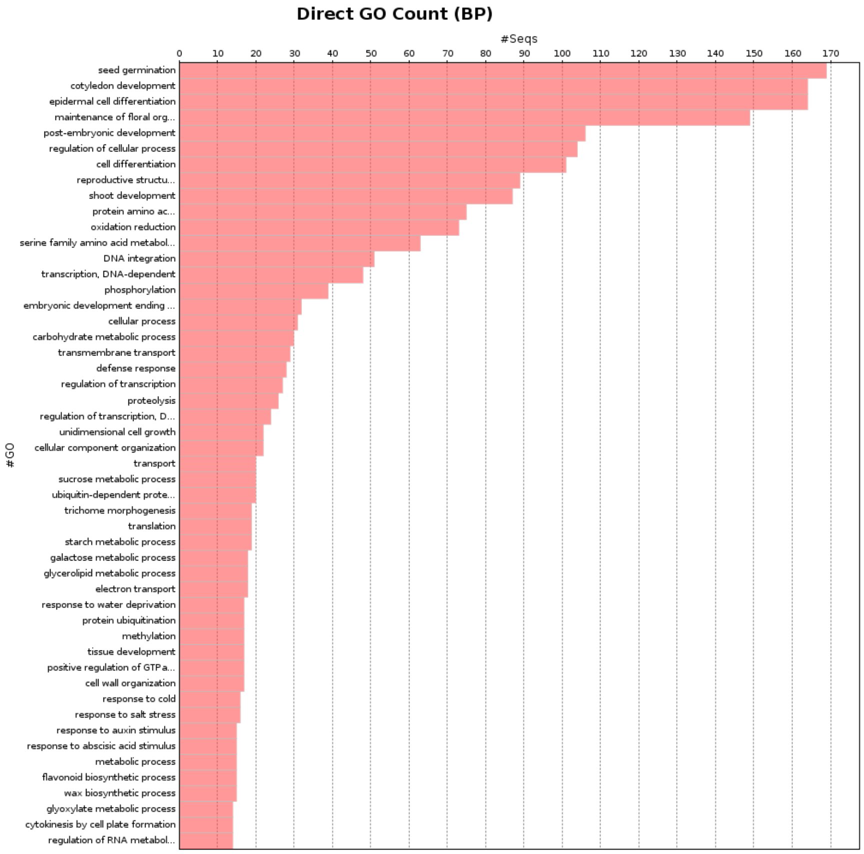


**Supplementary Figure S10**. Direct GO Count of the unique genes present in the BL1 genome annotated using Blast2go. Y-axis represents the GO categories and x-axis represents the number of unique genes in that category.

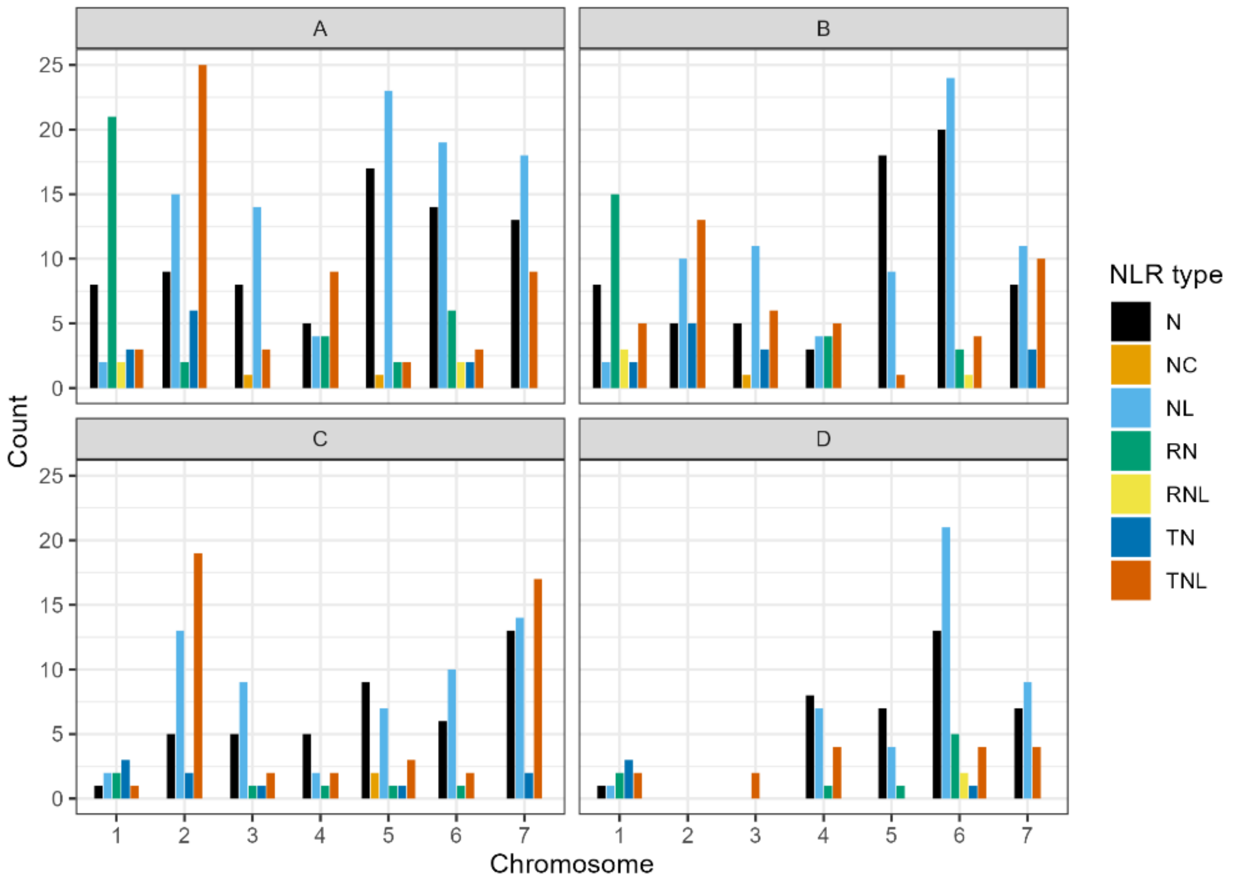


**Supplementary Figure S11.** Number of NLR genes in the BL1 genome containing the nucleotide-binding (NB) domains (NB-ARC). The domains are classified as follows: N = NB-ARC; NC = NB-ARC + CC (coiled coil); NL = NB-ARC + LRR (leucine-rich repeat); RN = RPW8 + NB-ARC; RNL = RPW8 + NB-ARC + LRR; TN = TIR (Toll-Interleukin 1 receptor) + NB-ARC; and TNL = TIR + NB-ARC + LRR.

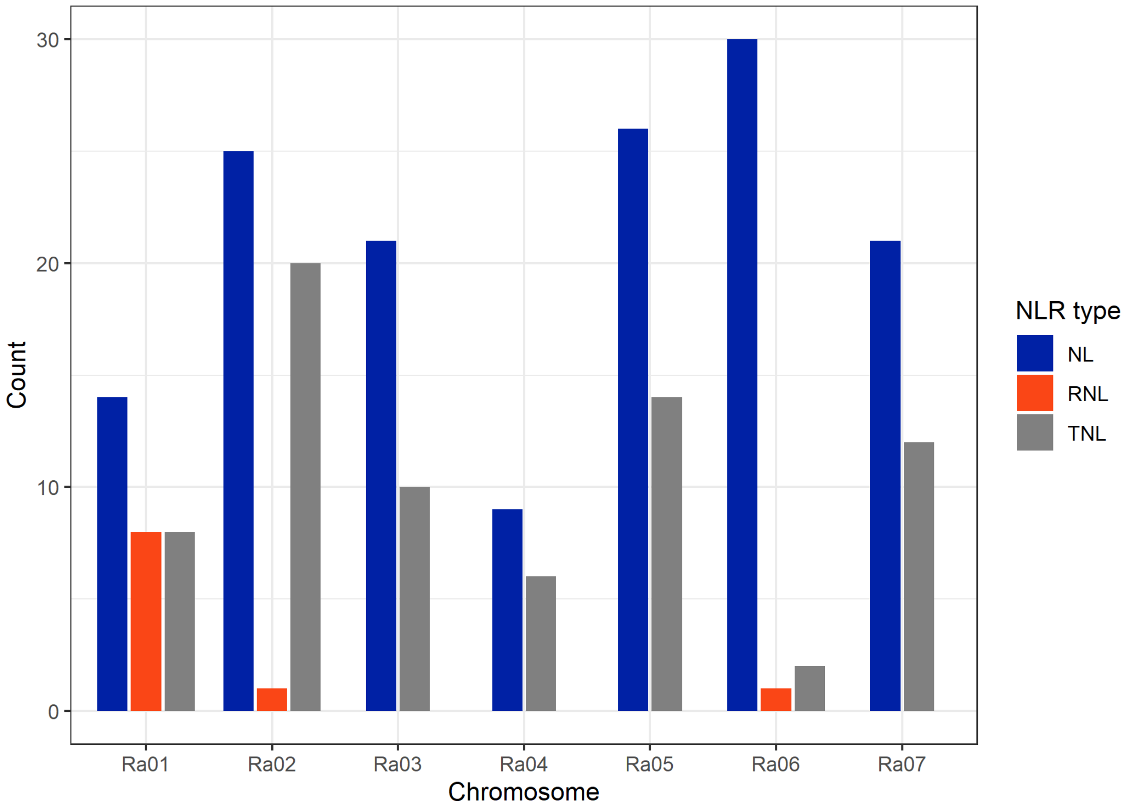


**Supplementary Figure S12.** Number of NLR genes in the Hillquist genome containing complete NBS domains. NLR types are classified as follows: NL = NB-ARC + LRR; RNL = RPW8 + NB-ARC + LRR; and TNL = TIR + NB-ARC + LRR.

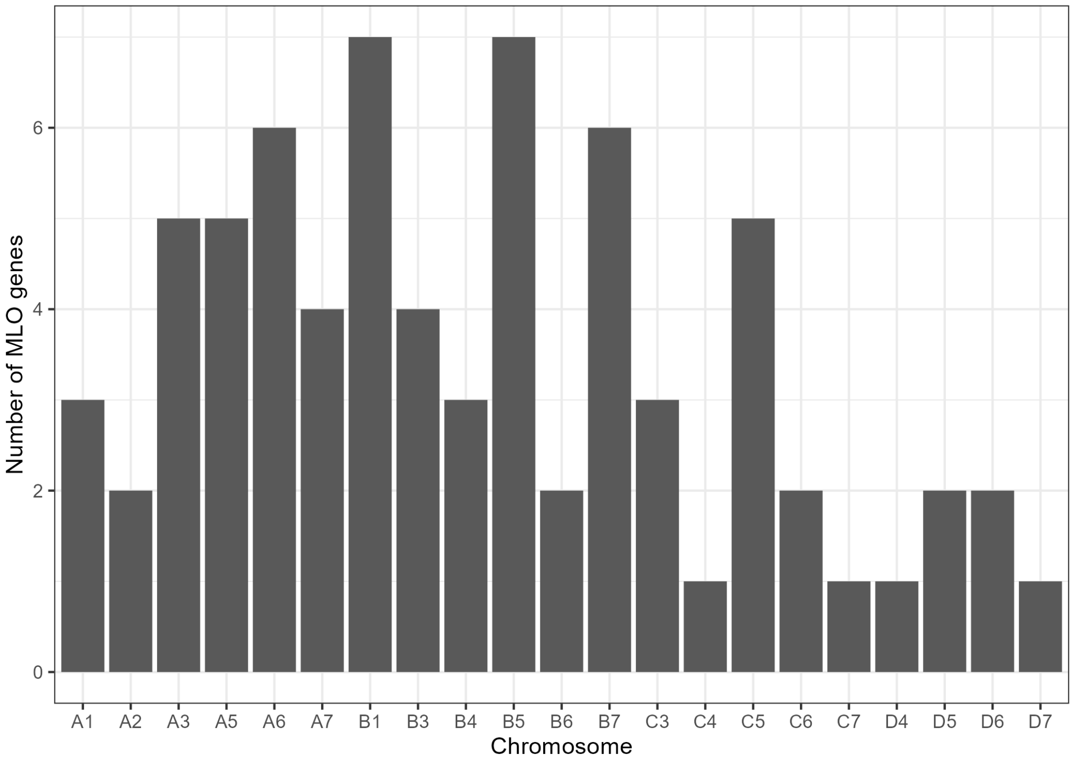


**Supplementary Figure S13.** Number of genes in the BL1 genome that are characterized as the *Mildew Locus O* (*MLO*) gene family members.

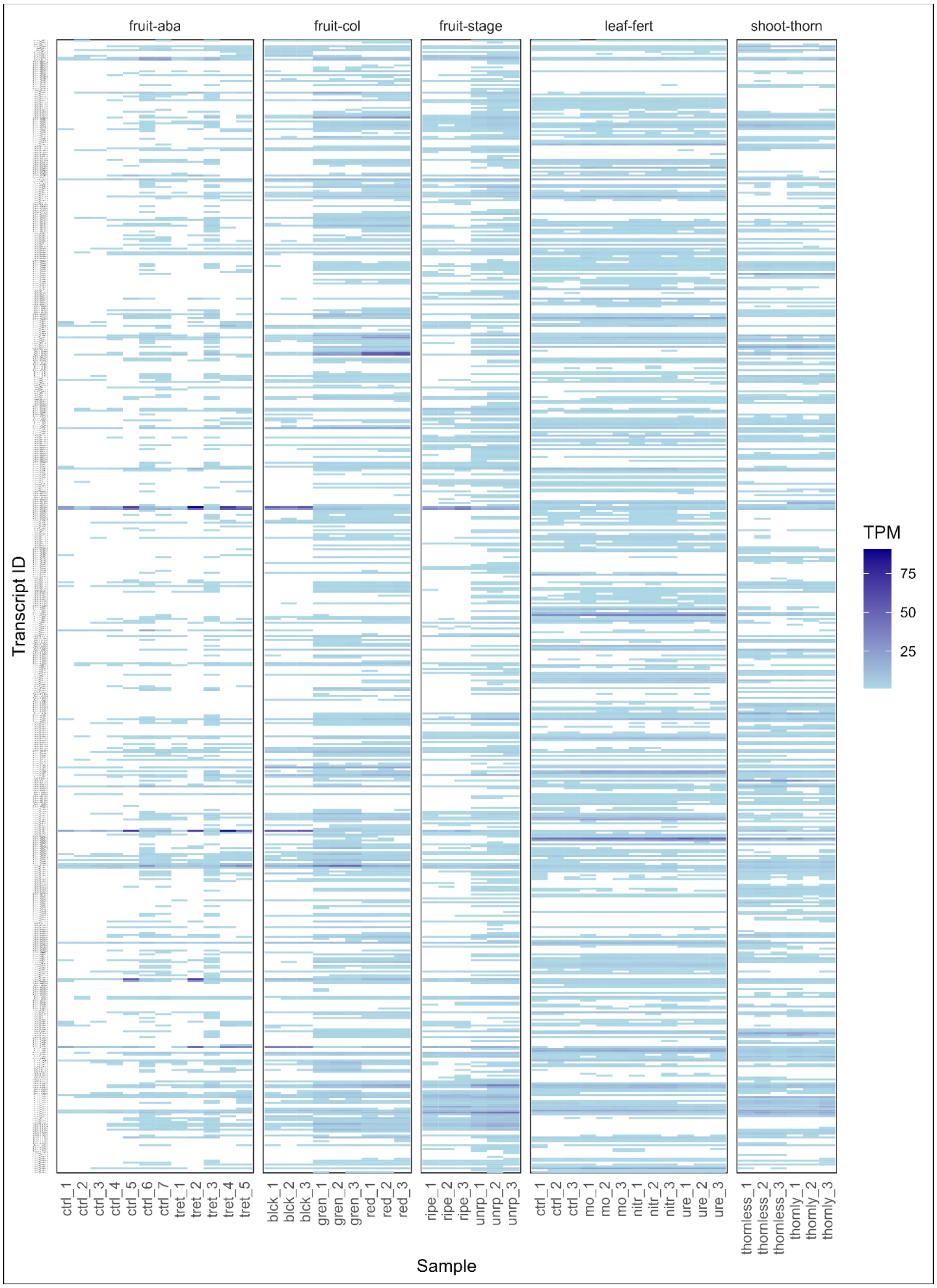


**Supplementary Figure S14.** Expression heatmap of the NLR genes expressed in different blackberry tissues. The normalized expression data as transcripts per million (TPM) for each transcript is shown. Five RNA-Seq experiments are shown: Fruit-aba (NCBI BioProject PRJNA680622), fruit-color (PRJNA744069), fruit-stage (PRJNA701162), leaf fertilizer (PRJNA787794), and shoot-thorn (data from this study).

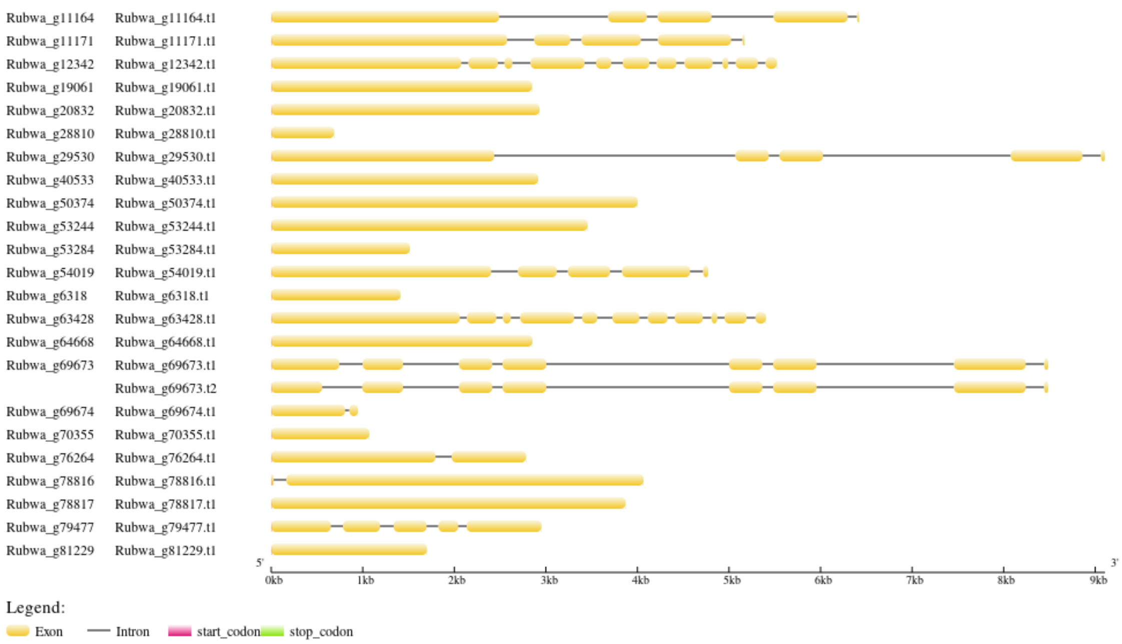


**Supplementary Figure S15.** Gene structure of commonly upregulated NLR transcripts. Two NLR transcripts (Rubwa_g19061.t1, Rubwa_g64668.t1) were commonly upregulated in both black and ripe berries while the remaining 22 NLR transcripts were commonly upregulated in green and unripe berries.

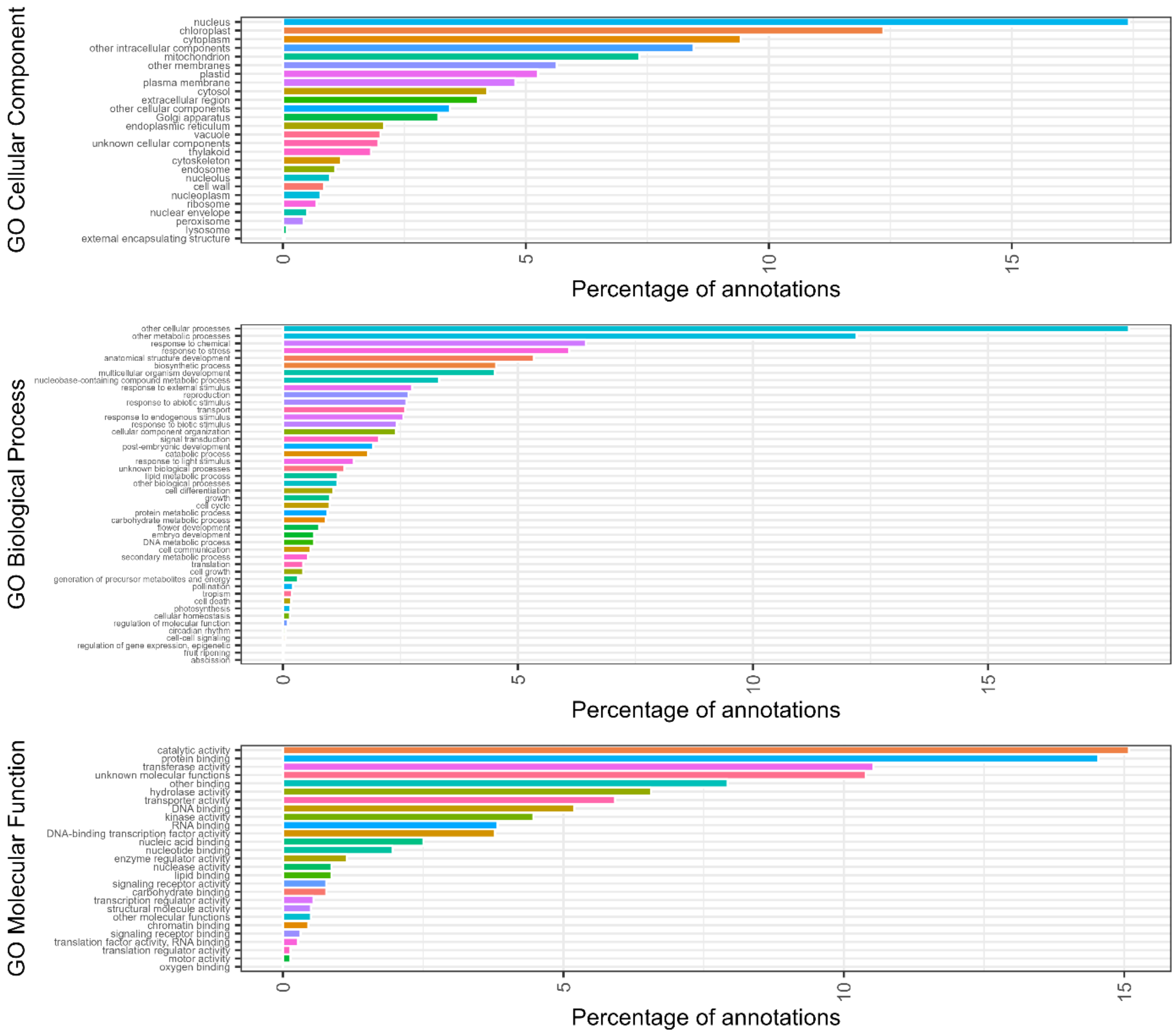


**Supplementary Figure S16.** GO characterization of genes present in the thorniness locus region.

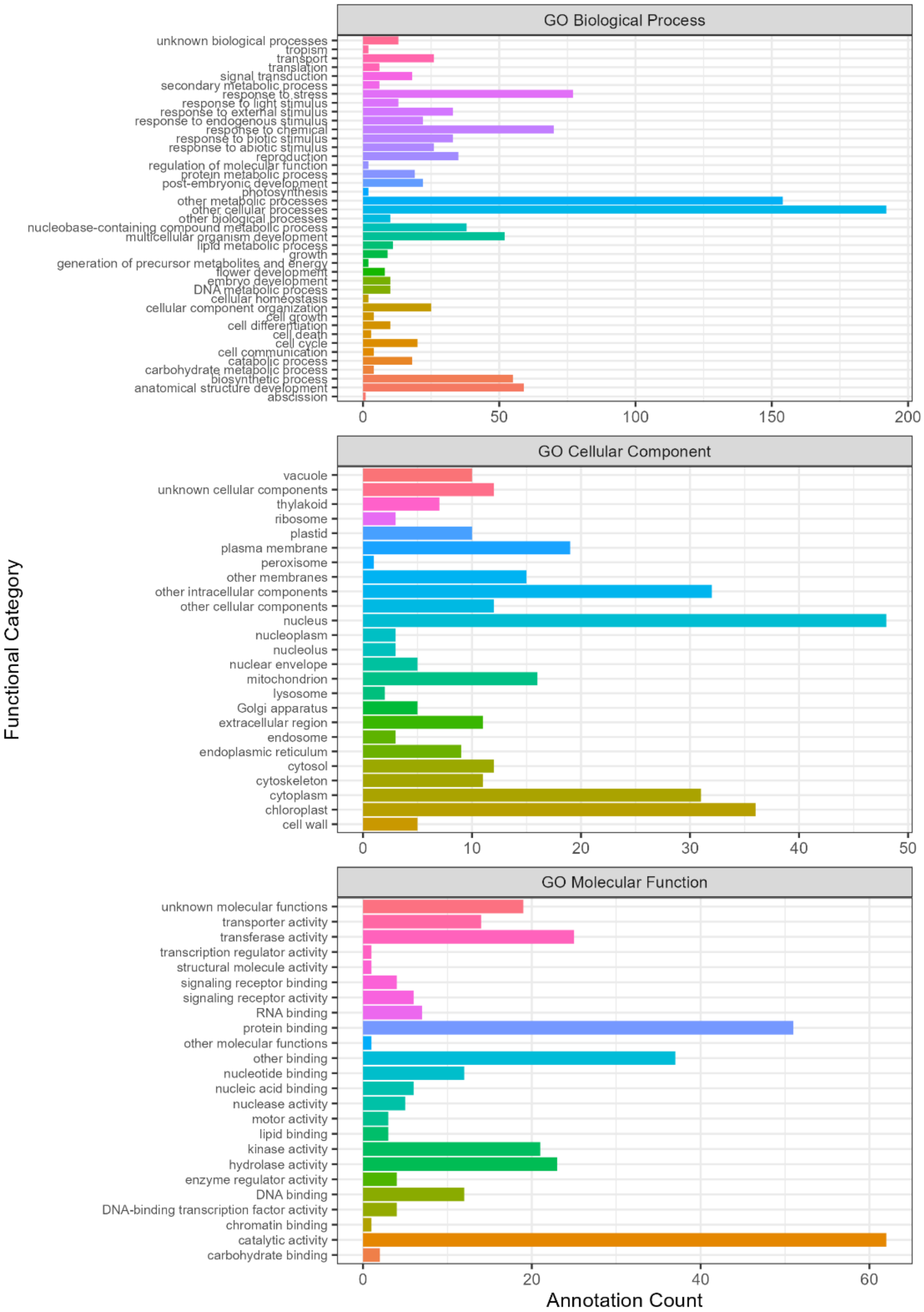


**Supplementary Figure S17.** GO characterization of genes in the thorniness locus region and with HIGH impact variants.

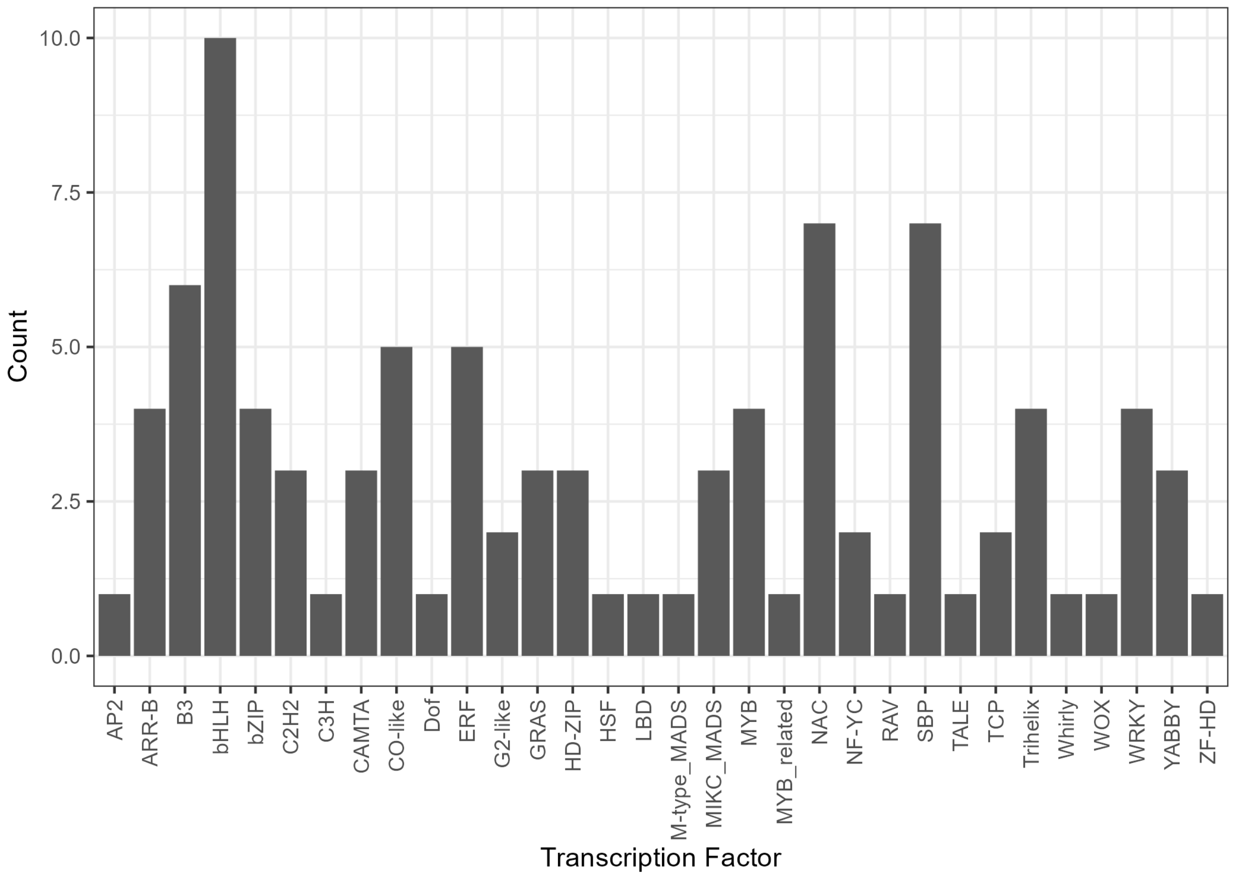


**Supplementary Figure S18.** Number of transcription factors encoded by genes in the thorniness locus region of BL1. X-axis represents the type of the transcription factor and y-axis represent the count of transcripts with the transcription factor. All transcripts present in the thorniness locus region were evaluated.
