## Supplementary Table for "A chromosome-scale and haplotype-resolved genome assembly of tetraploid blackberry (*Rubus* L. subgenus *Rubus* Watson)"

### Supplementary Tables

**Supplementary Table S1.** Summary of chromosome size, number of genes, GC content, number of NLR genes, and genetic loci present in each chromosome of the BL1 genome assembly.

| Name | Size (bp) | Telomers | Gene models (no) | GC content | NLR genes (no.) | Loci |
|---|---|---|---|---|---|---|
| A1 | 35,003,928 | 1 | 3,616 | 0.37 | 39 | |
| A2 | 45,097,959 | 1 | 5,355 | 0.37 | 58 | Primocane-fruiting |
| A3 | 37,827,979 | 1 | 3,398 | 0.37 | 29 | |
| A4 | 40,896,599 | 2 | 4,464 | 0.37 | 23 | Thorniness |
| A5 | 43,754,899 | 1 | 4,360 | 0.37 | 45 | |
| A6 | 55,812,704 | 1 | 6,903 | 0.37 | 47 | |
| A7 | 33,273,855 | 1 | 2,907 | 0.37 | 42 | |
| B1 | 38,009,712 | 0 | 3,782 | 0.37 | 35 | |
| B2 | 26,018,363 | 1 | 2,184 | 0.37 | 33 | |
| B3 | 43,292,644 | 1 | 3,931 | 0.37 | 27 | |
| B4 | 36,688,302 | 1 | 2,595 | 0.37 | 17 | |
| B5 | 33,267,598 | 1 | 2,426 | 0.37 | 29 | |
| B6 | 49,215,676 | 1 | 4,882 | 0.37 | 53 | |
| B7 | 35,734,842 | 2 | 2,984 | 0.37 | 34 | |
| C1 | 18,586,869 | 1 | 1,080 | 0.38 | 9 | |
| C2 | 30,546,578 | 0 | 2,313 | 0.38 | 40 | |
| C3 | 36,926,780 | 1 | 3,512 | 0.37 | 20 | |
| C4 | 21,383,817 | 0 | 1,449 | 0.37 | 10 | |
| C5 | 20,910,762 | 1 | 2,968 | 0.36 | 23 | |
| C6 | 34,137,567 | 0 | 3,024 | 0.37 | 23 | |
| C7 | 33,873,751 | 0 | 3,586 | 0.37 | 51 | |
| D1 | 17,890,200 | 0 | 1,058 | 0.37 | 9 | |
| D3 | 6,489,492 | 1 | 878 | 0.37 | 2 | |
| D4 | 30,667,472 | 1 | 2,375 | 0.37 | 20 | |
| D5 | 10,544,128 | 0 | 860 | 0.37 | 13 | |
| D6 | 35,448,538 | 1 | 3,587 | 0.37 | 46 | |
| D7 | 24,496,403 | 0 | 1,625 | 0.37 | 22 | |

**Supplementary Table S2.** BUSCO report of the BL1 genome assembly.

| Parameter | Count | Percentage (%) |
|---|---|---|
| Complete BUSCOs | 2,282 | 98.11 |
| Complete and single-copy BUSCOs | 622 | 26.74 |
| Complete and duplicated BUSCOs | 1,660 | 71.37 |
| Fragmented BUSCOs | 4 | 0.17 |
| Missing BUSCOs | 40 | 1.72 |
| **Total BUSCO groups searched** | 2,326 | |

**Supplementary Table S3.** Classification of repeat elements in the BL1 genome assembly.

| Class | Count | Length (bp) | Percentage (%) |
|---|---|---|---|
| SINEs | 1,520 | 118,787 | 0.01 |
| LINEs | 23,471 | 8,785,068 | 0.96 |
| LTR elements | 374,922 | 276,408,844 | 30.07 |
| DNA elements | 30,996 | 17,283,852 | 1.88 |
| Unclassified | 655,477 | 218,406,054 | 23.76 |
| Total interspersed repeats | | 521,002,605 | 56.68 |
| Small RNA | 2,981 | 2,069,051 | 0.23 |
| Satellites | 5,317 | 1,882,695 | 0.2 |
| Simple repeats | 200,431 | 8,824,431 | 0.96 |
| Low complexity | 34,180 | 1,875,785 | 0.2 |
| All repeat elements | | 535,654,567 | 58.27 |

**Supplementary Table S4**. Number of one-to-one orthologues shared between the BL1 genome and the fifteen public genomes in Rosaceae (woodland strawberry: *Fragaria vesca*, diploid blackberry: *Rubus argutus* cv. Hillquist, red raspberry: *Rubus idaeus* L. cv. Anitra, rose: *Rosa chinensis* 'Old Blush'; *R. chingii*; black raspberry: *R. occidentalis;* cherry; apple: *Malus domestica; P. avium; P. betulifolia; P. dulcis; P humilis; P. persica; P. sibirica; P. salicina*) and *Arabidopsis thaliana*.

(Attached Excel Sheet)

**Supplementary Table S5**. Summary of *MLO* genes identified in the BL1 genome.

(Attached Excel Sheet)

**Supplementary Table S6**. TPM values for NLR transcripts in different public and private blackberry RNA-Seq experiments.

(Attached Excel Sheet)

**Supplementary Table S7**. NLR transcripts commonly upregulated in two experiments. Transcriptome data for black and green berries were downloaded from NCBI BioProject PRJNA744069 (https://www.ncbi.nlm.nih.gov/bioproject/?term=PRJNA744069) and data for ripe and unripe berries were taken from PRJNA701162 (https://www.ncbi.nlm.nih.gov/bioproject/?term=PRJNA701162).

| Experiments | Chr | Start | End | Transcripts |
|---|---|---|---|---|
| Black and ripe | A5 | 15,147,893 | 15,150,742 | Rubwa_g19061.t1 |
| | C5 | 16,252,749 | 16,255,601 | Rubwa_g64668.t1 |
| Green and unripe | A2 | 30,216,297 | 30,217,709 | Rubwa_g6318.t1 |
| | A3 | 27,433,743 | 27,440,162 | Rubwa_g11164.t1 |
| | A3 | 27,471,880 | 27,477,048 | Rubwa_g11171.t1 |
| | A3 | 37,665,534 | 37,671,056 | Rubwa_g12342.t1 |
| | A5 | 40,197,234 | 40,200,164 | Rubwa_g20832.t1 |
| | A7 | 5,891,158 | 5,891,844 | Rubwa_g28810.t1 |
| | A7 | 14,576,714 | 14,585,820 | Rubwa_g29530.t1 |
| | B3 | 40,735,374 | 40,738,289 | Rubwa_g40533.t1 |
| | B6 | 45,992,833 | 45,996,834 | Rubwa_g50374.t1 |
| | B7 | 31,497,529 | 31,500,984 | Rubwa_g53244.t1 |
| | B7 | 32,003,610 | 32,005,124 | Rubwa_g53284.t1 |
| | C1 | 2,179,336 | 2,184,109 | Rubwa_g54019.t1 |
| | C5 | 7,191,407 | 7,196,809 | Rubwa_g63428.t1 |
| | C7 | 16,287,573 | 16,296,059 | Rubwa_g69673.t1 |
| | C7 | 16,287,573 | 16,296,059 | Rubwa_g69673.t2 |
| | C7 | 16,296,111 | 16,297,058 | Rubwa_g69674.t1 |
| | C7 | 25,353,492 | 25,354,562 | Rubwa_g70355.t1 |
| | D5 | 3,683,412 | 3,686,194 | Rubwa_g76264.t1 |
| | D6 | 25,136,256 | 25,140,321 | Rubwa_g78816.t1 |
| | D6 | 25,136,452 | 25,140,321 | Rubwa_g78817.t1 |
| | D6 | 29,887,379 | 29,890,329 | Rubwa_g79477.t1 |
| | D7 | 9,006,871 | 9,008,571 | Rubwa_g81229.t1 |

**Supplementary Table S8.** Summary of *DMR6* genes identified in the BL1 genome that are significantly up/down-regulated in different blackberry sample types.

(Attached Excel Sheet)

**Supplementary Table S9.** SnpEff summary of variants in the candidate genes located in the thorniness locus region that show HIGH impact.

(Attached Excel Sheet)

**Supplementary Table S10.** Summary of transcription factor genes located in the thorniness locus region.

| Transcript | TF | Arabidopsis TopHit | Pvalue | Function |
|---|---|---|---|---|
| Rubwa_g15974.t1 | RAV | AT1G25560.1 | 1.00E-153 | RAV family protein |
| Rubwa_g15008.t1 | bZIP | AT3G17609.2 | 2.00E-46 | HY5-homolog |
| Rubwa_g15055.t1 | SBP | AT3G60030.1 | 0 | squamosa promoter-binding protein-like 12 |
| Rubwa_g15056.t1 | SBP | AT3G60030.1 | 0 | squamosa promoter-binding protein-like 12 |
| Rubwa_g15075.t1 | CAMTA | AT5G09410.2 | 0 | ethylene induced calmodulin binding protein |
| Rubwa_g15099.t1 | Trihelix | AT2G33550.1 | 1.00E-104 | Trihelix family protein |
| Rubwa_g15184.t1 | Trihelix | AT2G33550.1 | 1.00E-104 | Trihelix family protein |
| Rubwa_g15200.t1 | bHLH | AT5G50915.2 | 5.00E-37 | bHLH family protein |
| Rubwa_g15209.t1 | bHLH | AT5G50915.2 | 6.00E-37 | bHLH family protein |
| Rubwa_g15220.t1 | NAC | AT5G13180.1 | 1.00E-105 | NAC domain containing protein 83 |
| Rubwa_g15240.t1 | NAC | AT5G13180.1 | 1.00E-105 | NAC domain containing protein 83 |
| Rubwa_g15253.t1 | SBP | AT5G43270.1 | 5.00E-73 | squamosa promoter binding protein-like 2 |
| Rubwa_g15266.t1 | CAMTA | AT1G67310.1 | 0 | Calmodulin-binding transcription activator protein with CG-1 and Ankyrin domains |
| Rubwa_g15270.t1 | SBP | AT5G50670.1 | 1.00E-45 | SBP family protein |
| Rubwa_g15277.t1 | WRKY | AT5G13080.1 | 5.00E-62 | WRKY DNA-binding protein 75 |
| Rubwa_g15306.t1 | NAC | AT5G13180.1 | 1.00E-105 | NAC domain containing protein 83 |
| Rubwa_g15328.t1 | bHLH | AT5G50915.2 | 5.00E-37 | bHLH family protein |
| Rubwa_g15334.t1 | CAMTA | AT1G67310.1 | 0 | Calmodulin-binding transcription activator protein with CG-1 and Ankyrin domains |
| Rubwa_g15344.t1 | SBP | AT5G43270.1 | 2.00E-84 | squamosa promoter binding protein-like 2 |
| Rubwa_g15361.t1 | NF-YC | AT5G43250.1 | 8.00E-44 | nuclear factor Y, subunit C13 |
| Rubwa_g15401.t1 | SBP | AT5G43270.1 | 2.00E-84 | squamosa promoter binding protein-like 2 |
| Rubwa_g15421.t1 | GRAS | AT1G14920.1 | 3.00E-76 | GRAS family protein |
| Rubwa_g15439.t1 | Trihelix | AT1G13450.1 | 0 | Trihelix family protein |
| Rubwa_g15522.t1 | NF-YC | AT5G43250.1 | 6.00E-43 | nuclear factor Y, subunit C13 |
| Rubwa_g15577.t1 | Trihelix | AT1G13450.1 | 0 | Trihelix family protein |
| Rubwa_g15596.t1 | GRAS | AT1G14920.1 | 1.00E-75 | GRAS family protein |
| Rubwa_g15837.t1 | bZIP | AT2G40950.1 | 1.00E-167 | bZIP family protein |
| Rubwa_g15874.t1 | bHLH | AT2G20180.1 | 6.00E-26 | phytochrome interacting factor 3-like 5 |
| Rubwa_g15906.t1 | bHLH | AT1G73830.1 | 8.00E-52 | BR enhanced expression 3 |
| Rubwa_g15927.t1 | LBD | AT3G02550.1 | 7.00E-67 | LOB domain-containing protein 41 |
| Rubwa_g15962.t1 | bZIP | AT1G68640.1 | 1.00E-175 | bZIP family protein |
| Rubwa_g15979.t1 | NAC | AT1G25580.1 | 1.00E-171 | NAC family protein |
| Rubwa_g15983.t1 | bHLH | AT1G68810.1 | 1.00E-116 | bHLH family protein |
| Rubwa_g15985.t1 | TCP | AT1G68800.1 | 6.00E-33 | TCP domain protein 12 |
| Rubwa_g16036.t1 | bZIP | AT3G30530.1 | 4.00E-49 | basic leucine-zipper 42 |
| Rubwa_g16044.t1 | bHLH | AT1G68920.3 | 1.00E-134 | bHLH family protein |

| Transcript | TF | Arabidopsis TopHit | Pvalue | Function |
|---|---|---|---|---|
| Rubwa_g16060.t1 | bHLH | AT1G69010.1 | 1.00E-114 | BES1-interacting Myc-like protein 2 |
| Rubwa_g16094.t1 | SBP | AT1G69170.1 | 2.00E-53 | SBP family protein |
| Rubwa_g16096.t1 | YABBY | AT1G69180.1 | 2.00E-61 | YABBY family protein |
| Rubwa_g16140.t1 | NAC | AT1G69490.1 | 3.00E-28 | NAC-like, activated by AP3/PI |
| Rubwa_g16146.t1 | WRKY | AT1G69310.2 | 7.00E-74 | WRKY DNA-binding protein 57 |
| Rubwa_g16176.t1 | WRKY | AT2G03340.1 | 1.00E-174 | WRKY DNA-binding protein 3 |
| Rubwa_g16208.t1 | NAC | AT1G69490.1 | 1.00E-120 | NAC-like, activated by AP3/PI |
| Rubwa_g16219.t1 | Dof | AT5G39660.1 | 1.00E-102 | cycling DOF factor 2 |
| Rubwa_g16233.t1 | ZF-HD | AT1G69600.1 | 1.00E-61 | zinc finger homeodomain 1 |
| Rubwa_g16237.t1 | NAC | AT1G26870.1 | 1.00E-103 | NAC family protein |
| Rubwa_g16249.t1 | TCP | AT3G47620.1 | 2.00E-62 | TEOSINTE BRANCHED, cycloidea and PCF (TCP) 14 |
| Rubwa_g16259.t1 | bHLH | AT1G26945.1 | 1.00E-39 | bHLH family protein |
| Rubwa_g16281.t1 | WRKY | AT5G15130.1 | 2.00E-83 | WRKY DNA-binding protein 72 |
| Rubwa_g16282.t1 | Whirly | AT1G14410.1 | 1.00E-131 | ssDNA-binding transcriptional regulator |
| Rubwa_g16392.t1 | YABBY | AT1G23420.1 | 3.00E-77 | YABBY family protein |
| Rubwa_g16392.t2 | YABBY | AT1G23420.1 | 7.00E-79 | YABBY family protein |
| Rubwa_g16432.t1 | HSF | AT1G67970.1 | 8.00E-88 | heat shock transcription factor A8 |
| Rubwa_g16521.t1 | GRAS | AT2G01570.1 | 0 | GRAS family protein |
| Rubwa_g16534.t1 | bHLH | AT1G10610.1 | 4.00E-69 | bHLH family protein |
| Rubwa_g15201.t1 | B3 | AT3G26790.1 | 6.00E-33 | B3 family protein |
| Rubwa_g15201.t2 | B3 | AT3G26790.1 | 8.00E-35 | B3 family protein |
| Rubwa_g15201.t3 | B3 | AT3G26790.1 | 7.00E-35 | B3 family protein |
| Rubwa_g15208.t1 | B3 | AT3G26790.1 | 8.00E-35 | B3 family protein |
| Rubwa_g15327.t1 | B3 | AT3G26790.1 | 8.00E-35 | B3 family protein |
| Rubwa_g15327.t2 | B3 | AT3G26790.1 | 7.00E-35 | B3 family protein |
| Rubwa_g16285.t1 | AP2 | AT3G54320.3 | 5.00E-91 | AP2 family protein |
| Rubwa_g15106.t1 | ERF | AT5G13330.1 | 2.00E-50 | related to AP2 6l |
| Rubwa_g15126.t1 | ERF | AT5G13910.1 | 7.00E-57 | ERF family protein |
| Rubwa_g15178.t1 | ERF | AT5G13330.1 | 2.00E-50 | related to AP2 6l |
| Rubwa_g15776.t1 | ERF | AT1G24590.1 | 3.00E-35 | DORNROSCHEN-like |
| Rubwa_g15935.t1 | ERF | AT1G68550.1 | 6.00E-40 | ERF family protein |
| Rubwa_g15227.t1 | CO-like | AT2G33500.1 | 1.00E-112 | B-box type zinc finger protein with CCT domain |
| Rubwa_g15247.t1 | CO-like | AT2G33500.1 | 1.00E-112 | B-box type zinc finger protein with CCT domain |
| Rubwa_g15314.t1 | CO-like | AT2G33500.1 | 1.00E-113 | B-box type zinc finger protein with CCT domain |
| Rubwa_g15857.t1 | CO-like | AT5G48250.1 | 2.00E-27 | B-box type zinc finger protein with CCT domain |
| Rubwa_g15928.t1 | CO-like | AT1G25440.1 | 5.00E-97 | B-box type zinc finger protein with CCT domain |
| Rubwa_g15416.t1 | ARR-B | AT1G67710.1 | 1.00E-109 | response regulator 11 |
| Rubwa_g15604.t1 | ARR-B | AT1G67710.1 | 1.00E-128 | response regulator 11 |
| Rubwa_g15859.t1 | ARR-B | AT4G16110.1 | 3.00E-34 | response regulator 2 |
| Rubwa_g16444.t1 | ARR-B | AT4G16110.1 | 1.00E-121 | response regulator 2 |
| Rubwa_g15971.t1 | G2-like | AT1G25550.1 | 1.00E-108 | G2-like family protein |

| Transcript | TF | Arabidopsis TopHit | Pvalue | Function |
|---|---|---|---|---|
| Rubwa_g16220.t1 | G2-like | AT1G69580.1 | 2.00E-91 | G2-like family protein |
| Rubwa_g15907.t1 | MIKC_MADS | AT2G22540.1 | 2.00E-70 | MIKC_MADS family protein |
| Rubwa_g16091.t1 | MIKC_MADS | AT1G69120.1 | 1.00E-112 | MIKC_MADS family protein |
| Rubwa_g16092.t1 | MIKC_MADS | AT2G03710.2 | 9.00E-89 | MIKC_MADS family protein |
| Rubwa_g16216.t1 | M-type_MADS | AT2G03060.2 | 1.00E-127 | AGAMOUS-like 30 |
| Rubwa_g16368.t2 | WOX | AT3G18010.1 | 8.00E-24 | WUSCHEL related homeobox 1 |
| Rubwa_g16262.t1 | HD-ZIP | AT1G69780.1 | 1.00E-122 | HD-ZIP family protein |
| Rubwa_g16298.t1 | HD-ZIP | AT3G01470.1 | 7.00E-41 | homeobox 1 |
| Rubwa_g16347.t1 | HD-ZIP | AT2G01430.1 | 6.00E-79 | homeobox-leucine zipper protein 17 |
| Rubwa_g15819.t1 | TALE | AT1G23380.1 | 1.00E-145 | KNOTTED1-like homeobox gene 6 |
| Rubwa_g15855.t1 | C3H | AT1G68200.1 | 1.00E-91 | C3H family protein |
| Rubwa_g16145.t1 | C2H2 | AT1G26610.1 | 5.00E-97 | C2H2-like zinc finger protein |
| Rubwa_g16412.t1 | C2H2 | AT1G24625.1 | 3.00E-41 | zinc finger protein 7 |
| Rubwa_g15905.t1 | MYB | AT1G68320.1 | 2.00E-90 | myb domain protein 62 |
| Rubwa_g16203.t1 | MYB | AT5G15310.1 | 1.00E-108 | myb domain protein 16 |
| Rubwa_g16217.t1 | MYB | AT1G69560.1 | 1.00E-77 | myb domain protein 105 |
| Rubwa_g16264.t1 | MYB | AT2G02820.2 | 0 | myb domain protein 88 |
| Rubwa_g16489.t1 | C2H2 | | | |
| Rubwa_g16144.t1 | MYB_related | | | |

**Supplementary Table S11.** Significantly regulated transcripts from the genes that are located in the thorniness locus region but changed expression in thornless blackberry samples.

| Regulation | Transcript | Chr | Start | End | Function |
|---|---|---|---|---|---|
| Down | Rubwa_g15061.t1 | A4 | 31,044,500 | 31,051,298 | Pleiotropic drug resistance 9 |
| Down | Rubwa_g15364.t1 | A4 | 32,837,052 | 32,838,794 | Caffeoyl-coa 3-O-methyltransferase |
| Down | Rubwa_g15365.t1 | A4 | 32,842,022 | 32,844,585 | Caffeoyl-coa 3-O-methyltransferase |
| Down | Rubwa_g15379.t1 | A4 | 32,916,431 | 32,919,922 | Leucine-rich repeat protein kinase family protein |
| Down | Rubwa_g15399.t1 | A4 | 33,043,356 | 33,045,350 | Glycosylphosphatidylinositol-anchored lipid protein transfer 1 |
| Down | Rubwa_g15426.t1 | A4 | 33,164,443 | 33,165,789 | HXXXD-type acyl-transferase family protein |
| Down | Rubwa_g15480.t1 | A4 | 33,392,345 | 33,394,461 | Cytochrome P450, family 716, subfamily A, polypeptide 1 |
| Down | Rubwa_g15517.t1 | A4 | 33,593,452 | 33,595,488 | Caffeoyl-coa 3-O-methyltransferase |
| Down | Rubwa_g15590.t1 | A4 | 34,059,719 | 34,061,011 | HXXXD-type acyl-transferase family protein |
| Down | Rubwa_g15695.t1 | A4 | 34,617,354 | 34,619,218 | Beta-ketoacyl reductase 1 |
| Down | Rubwa_g15696.t1 | A4 | 34,620,446 | 34,621,758 | Beta-ketoacyl reductase 1 |
| Down | Rubwa_g15776.t1 | A4 | 35,122,123 | 35,123,349 | DORNROSCHEN-like |
| Down | Rubwa_g15931.t2 | A4 | 35,921,666 | 35,923,361 | Homocysteine S-methyltransferase family protein |
| Down | Rubwa_g16559.t1 | A4 | 39,396,248 | 39,398,002 | Nine-cis-epoxycarotenoid dioxygenase 4 |
| Up | Rubwa_g15399.t2 | A4 | 33,043,371 | 33,045,350 | Glycosylphosphatidylinositol-anchored lipid protein transfer 1 |
| Up | Rubwa_g15453.t1 | A4 | 33,268,694 | 33,271,227 | Lysine histidine transporter 1 |
| Up | Rubwa_g15649.t1 | A4 | 34,377,053 | 34,380,182 | Mediator complex subunit Med28 |
| Up | Rubwa_g16203.t1 | A4 | 37,382,349 | 37,383,814 | MYB domain protein 16 |

**Supplementary Table S12.** Sequence of significantly up/down regulated transcripts in the thorniness locus region in the BL1 genome.

(Attached Excel Sheet)

**Supplementary Table S13.** Flowering related homologs identified in the primocane fruiting locus region in the BL1 genome.

(Attached Excel Sheet)

**Supplementary Table S14.** Summary of variants identified in seven blackberry cultivars/selections whose genes were re-sequenced in this study.

| Sample Name | PAF | PA45m | BL2 | Traveler | PA45 | Kiowa | Osage |
|---|---|---|---|---|---|---|---|
| SNPs | 1,770,525 | 3,250,681 | 3,572,942 | 1,919,694 | 3,288,733 | 4,526,044 | 4,375,660 |
| MNPs | 250,516 | 506,824 | 497,003 | 260,270 | 517,574 | 681,958 | 682,537 |
| Insertions | 102,917 | 173,458 | 192,696 | 111,710 | 175,908 | 234,679 | 229,344 |
| Deletions | 112,886 | 192,806 | 218,002 | 125,626 | 196,740 | 260,120 | 254,018 |
| Indels | 82,477 | 156,045 | 163,141 | 81,311 | 159,661 | 211,092 | 211,775 |
| Same as reference | 7,433,135 | 5,520,894 | 4,476,908 | 7,509,798 | 5,491,356 | 3,701,194 | 4,247,314 |
| Missing Genotype | 2,275,136 | 2,226,884 | 2,906,900 | 2,019,183 | 2,197,620 | 2,412,505 | 2,026,944 |
| SNP Transitions/Transversions | 1.74 (1286924/ 738419) | 1.75 (2545018/ 1455536) | 1.79 (2996469/ 1674758) | 1.80 (1485331/ 823946) | 1.75 (2580056/ 1474901) | 1.77 (3650832/ 2061786) | 1.75 (3530449/ 2022724) |
| Total Het/Hom ratio | 6.12 (1993545/ 325776) | 3.42 (3311176/ 968638) | 2.31 (3242666/ 1401118) | 4.12 (2010344/ 488267) | 3.37 (3344982/ 993634) | 2.92 (4406046/ 1507847) | 2.81 (4244116/ 1509218) |
| SNP Het/Hom ratio | 5.99 (1517255/ 253270) | 3.38 (2508180/ 742501) | 2.29 (2485639/ 1087303) | 3.95 (1532126/ 387568) | 3.33 (2529937/ 758796) | 2.87 (3355742/ 1170302) | 2.76 (3212061/ 1163599) |
| MNP Het/Hom ratio | 6.62 (217645/ 32871) | 3.83 (401791/ 105033) | 2.63 (359968/ 137035) | 5.04 (217208/ 43062) | 3.73 (408052/ 109522) | 3.49 (530148/ 151810) | 3.40 (527436/ 155101) |
| Insertion Het/Hom ratio | 6.33 (88886/ 14031) | 3.28 (132951/ 40507) | 2.20 (132507/ 60189) | 4.40 (91015/ 20695) | 3.21 (134129/ 41779) | 2.72 (171516/ 63163) | 2.56 (164960/ 64384) |
| Deletion Het/Hom ratio | 6.10 (96986/ 15900) | 3.00 (144654/ 48152) | 2.07 (147049/ 70953) | 4.31 (101981/ 23645) | 2.97 (147189/ 49551) | 2.50 (185802/ 74318) | 2.36 (178338/ 75680) |
| Indel Het/Hom ratio | 7.50 (72773/ 9704) | 3.81 (123600/ 32445) | 2.57 (117503/ 45638) | 5.11 (68014/ 13297) | 3.70 (125675/ 33986) | 3.37 (162838/ 48254) | 3.20 (161321/ 50454) |
| Insertion/Deletion ratio | 0.91 (102917/ 112886) | 0.90 (173458/ 192806) | 0.88 (192696/ 218002) | 0.89 (111710/ 125626) | 0.89 (175908/ 196740) | 0.90 (234679/ 260120) | 0.90 (229344/ 254018) |
| Indel/SNP+MNP ratio | 0.15 (298280/ 2021041) | 0.14 (522309/ 3757505) | 0.14 (573839/ 4069945) | 0.15 (318647/ 2179964) | 0.14 (532309/ 3806307) | 0.14 (705891/ 5208002) | 0.14 (695137/ 5058197) |

**Supplementary Table S15.** Variants located within the NLR genes between leaf rust-susceptible and resistant blackberry cultivars/selections.

(Attached Excel Sheet)

**Supplementary Table S16.** Main characteristics of blackberry cultivars and selections whose genomes were sequenced or re-sequenced in this study.

| **Blackberry cultivars/selections** | **Thorns** | **Fruiting habit** | **Susceptibility to leaf spot caused by *Pseuodocercospora pancratii* (Marin et al., 2023)** | **Resistance to leaf rust caused by *Kuehneola uredines* (Marin et al., 2022)** |
|---|---|---|---|---|
| BL1 | No | Primocane | Unknown | Unknown |
| BL2 | No | Primocane | Unknown | Unknown |
| Kiowa | Yes | Floricane | Highly susceptible | Susceptible |
| Osage | No | Floricane | Susceptible | Resistant |
| Prime-Ark® 45 (PA45) | Yes | Primocane | Unknown | Resistant |
| Prime-Ark® 45 variant (PA45m) | Yes, fewer, shorter | Primocane | Unknown | Unknown |
| Prime-Ark® Freedom (PAF) | No | Primocane | Highly susceptible | Susceptible |
| Prime-Ark® Traveler (PAT) | No | Primocane | Susceptible | Resistant |

**Supplementary Table S17.** Results of different genome assemblers for initial assembly.

| Feature / Software | CANU | wtdbg2 | WENGAN | smartdenovo | necat | flye |
|---|---|---|---|---|---|---|
| **Contig number** | 5,616 | 7,914 | 5,397 | 2,310 | 1,037 | 9,064 |
| **Total length** | 1,078,629,771 | 575,938,167 | 293,747,100 | 813,974,517 | 875,604,717 | 796,106,916 |
| **Max length** | 9,724,148 | 8,372,833 | 1,212,355 | 5,832,673 | 18,743,802 | 6,541,860 |
| **N50** | 453,091 | 202,570 | 105,074 | 634,739 | 2,220,393 | 262,817 |
| **N90** | 69,686 | 28,604 | 24,369 | 162,109 | 431,081 | 47,261 |

**Supplementary Table S18.** Parameter evaluation for NECAT assembly.

| Parameter | Default | Setting1 | Setting2 | Setting3 | Setting4 | Setting5 | Setting6 |
|---|---|---|---|---|---|---|---|
| **Minimum read length** | 3,000 | 5000 | 5,000 | 5,000 | 3,000 | 3,000 | 3,000 |
| **Prep output coverage** | 40 | 40 | 50 | 60 | 40 | 60 | 60 |
| **Consensus output coverage** | 30 | 30 | 40 | 50 | 20 | 40 | 30 |
| **Sequence** | 1,037 | 1,018 | 1,735 | 1,320 | 1,672 | 1,252 | 727 |
| **Total length** | 875,604,717 | 859,223,131 | 886,143,963 | 914,686,082 | 908,021,257 | 908,624,256 | 782,234,984 |
| **Max length** | 18,743,802 | 17,931,999 | 18,307,468 | 13,723,711 | 9,724,569 | 15,579,035 | 16,003,244 |
| **N50** | 2,220,393 | 2,098,452 | 1,531,495 | 1,806,665 | 1,217,099 | 1,967,596 | 1,952,295 |
| **N90** | 431,081 | 423,350 | 241,112 | 341,438 | 254,024 | 356,952 | 439,839 |
